# PTRAMP, CSS and Ripr form a conserved complex required for merozoite invasion of *Plasmodium* species into erythrocytes

**DOI:** 10.1101/2025.03.25.644866

**Authors:** Benjamin A. Seager, Pailene S. Lim, Keng Heng Lai, Lionel Brice Feufack-Donfack, Sheena Dass, Xiao Xiao, Nicolai C. Jung, Anju Abraham, Matthew J. Grigg, Nicholas M. Anstey, Timothy William, Jetsumon Sattabongkot, Andrew Leis, Rhea J. Longley, Manoj T. Duraisingh, Jean Popovici, Danny W. Wilson, Stephen W. Scally, Alan F. Cowman

## Abstract

Invasion of erythrocytes by members of the *Plasmodium* genus is an essential step of the parasite lifecycle, orchestrated by numerous host-parasite interactions. In *P. falciparum* Rh5, with PfCyRPA, PfRipr, PfCSS, and PfPTRAMP, forms the essential PCRCR complex which binds basigin on the erythrocyte surface. Rh5 is restricted to *P. falciparum* and its close relatives; however, PTRAMP, CSS and Ripr orthologs are present across the *Plasmodium* genus. We investigated PTRAMP, CSS and Ripr orthologs from three species to elucidate common features of the complex. Like *P. falciparum,* PTRAMP and CSS form a disulfide-linked heterodimer in both *P. vivax* and *P. knowlesi* with all three species forming a complex (PCR) with Ripr by binding its C-terminal region. Cross-reactive antibodies targeting the PCR complex differentially inhibit merozoite invasion. Cryo-EM visualization of the *P. knowlesi* PCR complex confirmed predicted models and revealed a core invasion scaffold in *Plasmodium* spp. with implications for vaccines targeting multiple species of malaria-causing parasites.

## Introduction

There are more than 200 species of *Plasmodium* that infect a diverse range of hosts including primates, rodents, reptiles and birds. At least six species, including *Plasmodium falciparum, P. vivax* and *P. knowlesi,* have the ability to infect humans. *P. falciparum* is the most lethal species to infect humans, while *P. vivax* is the most widespread globally^1,2^. *P. knowlesi* is confined mainly to regions of Southeast Asia and is transmitted from macaques by zoonotic infection^3,4^. The *Plasmodium* genus can be divided into three main subgenera or clades that include *Laverania* (includes *P. falciparum*), *Plasmodium* (includes most human infective species such as *P. vivax* and *P. knowlesi*) and *Vinckeia* (primarily rodent infective species).

The three clades of the *Plasmodium* genus represent distinct evolutionary branches distinguished by geographic distribution and severity of disease; however, they can also exhibit distinct host cell selectivity. *P. vivax* has a strict preference for invasion of reticulocytes in the blood whereas *P. falciparum* can invade both reticulocytes and the more mature normocytes^5^. While the core machinery for invasion of reticulocytes and normocytes by *P. vivax* and *P. falciparum*, such as the parasite actomyosin motor, is conserved, there are distinct ligand-receptor interactions that provide selectivity for host cell invasion (reviewed in ^6^). In *P. falciparum,* many of these ligands are dispensable for invasion^7^. The notable exception is reticulocyte-binding protein homologue 5 (Rh5)^8,9^ which is an essential *P. falciparum* ligand that binds to the receptor basigin on human erythrocytes^10^. Rh5 can play a role in host tropism through polymorphisms in the protein and differential binding to basigin of other non-human primates^8,11^.

In *P. falciparum* Rh5 functions in a complex of five proteins that include CyRPA (Cysteine Rich Protective Antigen)^12^, Ripr (Rh5 interacting protein)^13^, PTRAMP (*Plasmodium* thrombospondin-related apical merozoite protein)^14,15^, and CSS (cysteine-rich, small, secreted)^16^ that has been termed the PCRCR complex^17–19^. PTRAMP and CSS form a disulfide-linked heterodimer that tethers the PCRCR complex to the merozoite membrane via the transmembrane domain of PTRAMP^18^. All proteins in the PCRCR complex are functionally essential for *P. falciparum* merozoite invasion of human erythrocytes and, crucially, antibodies and nanobodies that bind to individual proteins can inhibit merozoite invasion^18,20–25^. Conditional gene knockouts of each PCRCR protein in *P. falciparum* display the same phenotype whereby the merozoites can interact with the erythrocyte surface and produce the strong deformation of the host membrane typical during normal invasion; however, the merozoite fails to internalize^18,26^. The PCRCR complex has been hypothesized to capture and anchor the increased membrane surface contact formed between the merozoite and erythrocyte membrane that is created during strong deformation driven by the merozoites actomyosin motor^18^. This facilitates the establishment of the moving junction and is followed by the downstream events of invasion and ultimately the internalization of the merozoite into the erythrocyte^27,28^.

Despite being essential for *P. falciparum* invasion, Rh5 orthologs are absent in species outside of the *Laverania* subgenus^29,30^. Consequently, utilization of basigin as a host receptor for invasion is not universal, as demonstrated for both *P. knowlesi* and *P. vivax*^16^, suggesting that other parasite ligand-receptor interactions facilitate host cell attachment in other *Plasmodium* spp. Non-*Laverania* species do, however, possess homologues of other components of the *P. falciparum* PCRCR complex and *P. knowlesi* orthologs of PTRAMP, CSS and Ripr have been shown to be essential for merozoite invasion using conditional gene knockouts^16^ and more recently a high-resolution transposon mutagenesis screen for growth^31,32^. It has been suggested that PkPTRAMP, PkCSS and PkRipr form a complex and that PkPTRAMP provides the means for erythrocyte binding^16^. The PkCyRPA homologue was also identified, and while it has been shown to be essential for parasite growth, it does not appear to be part of this complex in *P. knowlesi*^16^.

Here we hypothesized that PTRAMP, CSS and Ripr form a common basis for invasion complexes across *Plasmodium* spp. We leveraged recent insights into the PCRCR complex to characterize these proteins in several species of *Plasmodium* to elucidate the conserved features of merozoite invasion complexes. Our findings revealed a conserved PCR trimeric complex common to all *Plasmodium* clades that forms a core invasion scaffold. Additionally, cross-reactive antibodies targeting PTRAMP, CSS, and Ripr were identified that differentially inhibit merozoite invasion of erythrocytes, including an antibody targeting Ripr that inhibited both *P. knowlesi* and *P. falciparum* growth. Identification of a conserved molecular scaffold presents an attractive approach for the development of vaccines targeting multiple species of malaria-causing parasites.

## Results

### PTRAMP, CSS and Ripr are conserved in all clades of *Plasmodium*

We first confirmed that the proteins constituting the *P. falciparum* PCRCR complex are conserved in other *Plasmodium* spp. by searching for orthologs and found that PTRAMP, CSS and Ripr were present in all subgenera **(Fig. 1a)**. This conservation contrasts with Rh5 which is only present in *P. falciparum* and other *Laverania* species. Whilst CyRPA is relatively conserved, none of the species within the *Vinckeia* subgenus possess a CyRPA ortholog. This suggests PTRAMP, CSS and Ripr (PCR) form a conserved three-membered complex present across all *Plasmodium* spp.

**Fig. 1.**
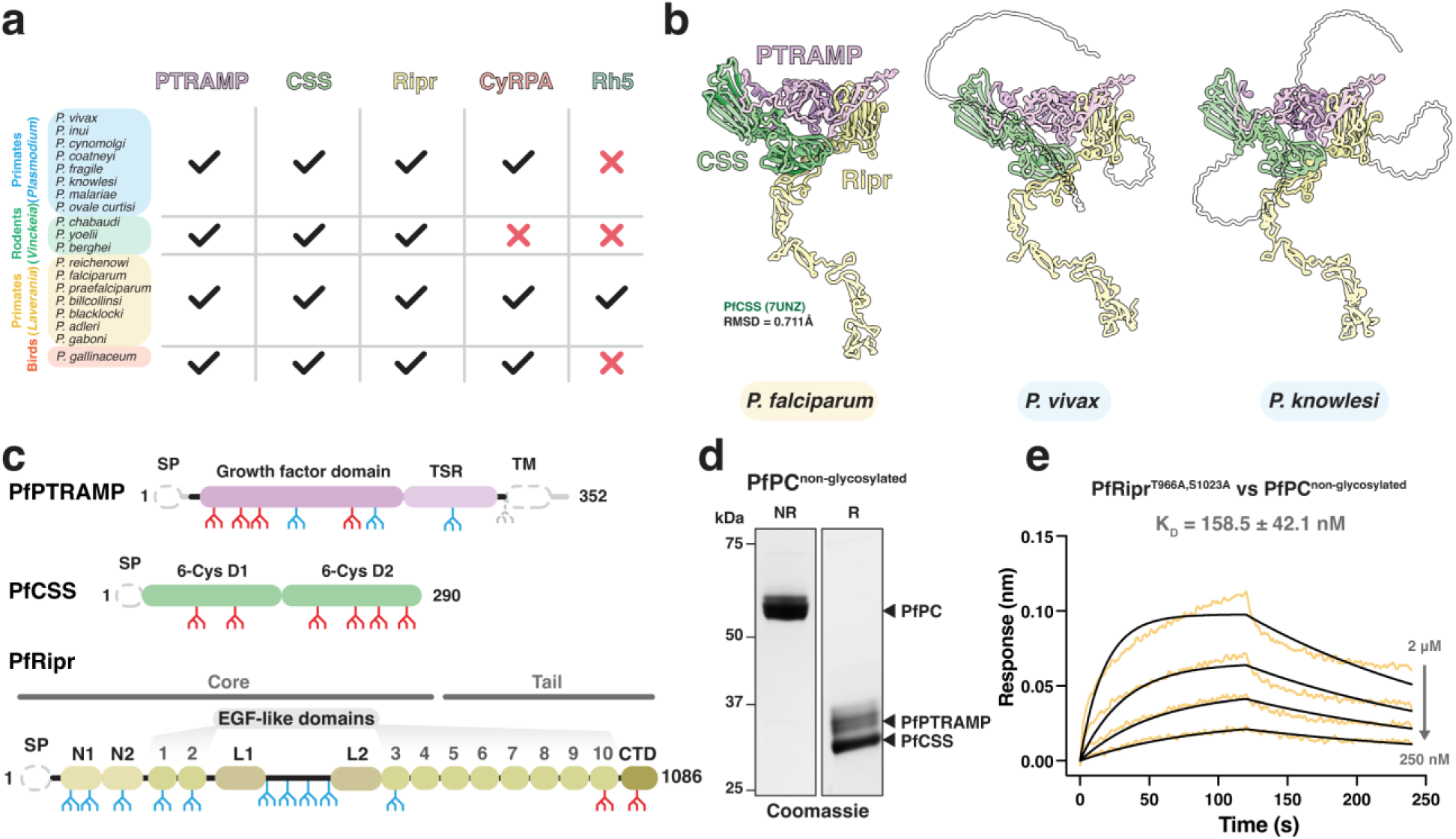
PTRAMP, CSS, and Ripr are common to all clades of *Plasmodium*. **a.** Comparison of PCRCR orthologs in the *Plasmodium* genus, showing that PTRAMP, CSS, and Ripr are common to all species, whereas Rh5 is restricted to the *Laverania* subgenus, and CyRPA is absent from the rodent-infective species (*Vinckeia*). **b.** AlphaFold 3 predictions of PTRAMP, CSS, and the Ripr tail show a conserved architecture is predicted for each species. For the *P. falciparum* predicted complex, the previously published structure for PfCSS is superimposed in dark green. Regions that are predicted to be disordered, and therefore have low model confidence, are shown as transparent. The transmembrane domain and signal peptide have been removed from PTRAMP and CSS for clarity **c.** Domain diagrams for PfPTRAMP, PfCSS and PfRipr showing the predicted N-linked glycosylation sites^35^ (blue) that were either mutated (red) or the sequon truncated (grey). Grey regions indicate the stretches of sequence either processed (signal peptides) or removed for recombinant expression, in the case of PfPTRAMP (transmembrane domain and cytoplasmic tail). **d.** SDS-PAGE of non-glycosylated PfPC in both non-reduced (NR) and reduced (R) conditions. **e.** A representative biolayer interferometry sensorgram of non-glycosylated PfRipr binding to non-glycosylated PfPC showing data (yellow) and 1:1 model best fit (black).

AlphaFold 3 was used to predict the structure of the PCR complex of different *Plasmodium* spp. to understand the assembly of the three proteins **(Fig. 1b, Extended data Fig. 1)**^33,34^. This revealed a common architecture of the PCR complex in which PTRAMP and CSS together form a platform that is bound by the C-terminal end of Ripr. The PfCSS crystal structure aligned with the AlphaFold 3 prediction of PfPCR with an RMSD of 0.711 Å, suggesting that no conformational changes are required within PfCSS to facilitate PfRipr binding **(Fig. 1b)**.

The predicted engagement of Ripr with both PTRAMP and CSS in the AlphaFold 3 models was similar in all *Plasmodium* spp. but was inconsistent with existing biophysical data for *P. falciparum* which had suggested that PfCSS alone was sufficient for PfRipr binding^18^. However, this interaction was low affinity with K_D_ of CSS binding to Ripr in the low micromolar range^18,19^. In contrast, the PCR model predicted significant interaction of both PTRAMP and CSS with Ripr for *P. falciparum*, *P. vivax* and *P. knowlesi*. PfPTRAMP, PfCSS and PfRipr all contain multiple predicted N-linked glycan motifs and whilst there is very limited glycosylation in *P. falciparum* these proteins were expressed in insect cells and therefore are predicted to undergo extensive glycosylation **(Fig 1c)**^35^. To further investigate PfPTRAMP-PfCSS (PfPC) - PfRipr binding, we recombinantly expressed glycosylation-modified variants that either completely lacked glycans or had significantly reduced glycosylation **(Fig 1c, d)**.

Using biolayer interferometry (BLI), we determined that non-glycosylated forms of PfRipr and PfPC interacted with a K_D_ of 160 nM, representing a ∼10-fold stronger interaction compared to previous measurements (1-4 μM)^18,19^ and equivalent to the affinity measured between Rh5 and CyRPA (179 nM) **(Fig. 1e)**^36^. The enhanced binding affinity of non-glycosylated proteins suggests that N-linked glycans on PfRipr and PfPC, added during heterologous expression, had interfered with their interaction interface in previous studies. Despite removal of a majority of the PfPTRAMP glycans monomeric PfPTRAMP showed no binding to PfRipr **(Supp. Fig. 1)**. These results suggest that the complete PfPC heterodimer is necessary for high-affinity PfRipr binding.

### The *P. vivax* orthologs of PTRAMP and CSS form a disulfide-linked heterodimer

To investigate whether the disulfide-linked PC heterodimer is conserved as the basis for Ripr binding across *Plasmodium* species, as predicted by AlphaFold 3^34^, we co-expressed *P. vivax* PTRAMP and CSS orthologs in mammalian cells. The resulting PvPC heterodimer could be separated into its component monomers through reduction of the intermolecular disulfide bond **(Fig. 2a)**. Nanobodies were raised against the purified PvPC heterodimer to enable further structural and biophysical characterization **(Supp. Fig. 2)**. Crystallization of the PvPC heterodimer was achieved by truncating the predicted disordered N-terminal repeat region of PvCSS and adding nanobody D7 **(Fig. 2b, Table S1)**. The resulting structure confirmed that PvCSS adopts the previously characterized two-domain degenerate 6-Cys fold seen in PfCSS **(Fig. 2c, Extended data Fig. 2)**^18^.

**Fig. 2.**
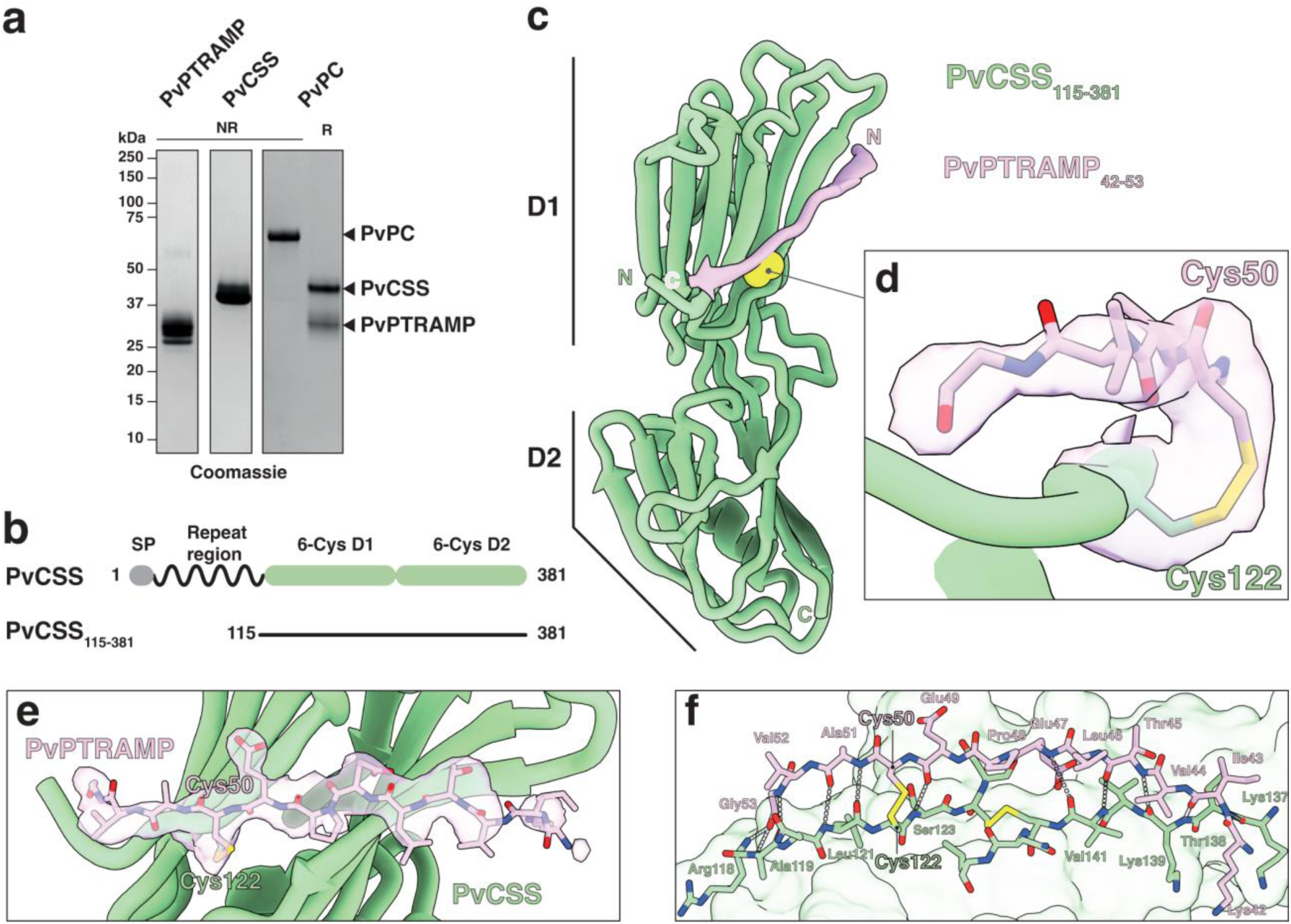
Structure of the intermolecular disulfide bond between PvCSS and PvPTRAMP. **a.** SDS-PAGE of recombinantly expressed PvPTRAMP, PvCSS and PvPC heterodimer. **b.** Domain diagram of PvCSS showing the disordered repeat region at the N-terminus. **c.** Crystal structure of PvCSS_115-381_(green) and PvPTRAMP_42-53_(pink) with the disulfide formed between them in space-filling atomic depiction (yellow). **d.** Detail of the intermolecular disulfide bond between PvCSS and PvPTRAMP. Density is contoured at 1.0 σ and density extends to a range of 1.8 Å. **e.** A model of PvCSS and PvPTRAMP_42-53_, showing the region of the unbiased electron density omit map that was attributed to PvPTRAMP. Density is represented as in d). **f.** The interface between PvPTRAMP_42-53_ and PvCSS. All intermolecular hydrogen bonds formed are represented in grey.

No electron density was observed for either the growth-factor domain (GFD) or the thrombospondin repeat (TSR) domain of PvPTRAMP, despite space being available within the crystal lattice **(Extended data Fig. 2c)**. This suggests that the majority of PvPTRAMP was insufficiently stabilized within the crystal lattice to produce coherent diffraction. Nevertheless, clear electron density extended from PvCSS cysteine 122, the predicted site of disulfide formation with PvPTRAMP **(Fig. 2d)**. Modelling of PvPTRAMP residues 42 to 53, revealed the structural basis for PvPC heterodimerization **(Fig. 2e, f)**. Specifically, PvPTRAMP forms an interrupted β- strand that extends across both β-sheets of the CSS D1 domain, establishing multiple backbone interactions **(Fig. 2c, f, Table S2)**.

Alignment of available PTRAMP and CSS sequences showed that the two cysteines involved in heterodimerization are conserved across most species **(Supp. Fig. 3, 4)**. One exception is *Plasmodium inui*, which has tyrosine and serine substitutions in PTRAMP and CSS, respectively **(Supp. Fig. 3, 4)**. Nevertheless, AlphaFold^33^ modelling predicts a similar interface between PTRAMP and CSS in this region **(Extended data Fig. 1d)**.

### PvRipr binds the PvPC heterodimer to form a high affinity complex

Biophysical analysis revealed that PvPC binds to PvRipr with high affinity (K_D_ = 28.8 ± 3.9 nM) **(Fig. 3a, Extended data Fig. 3a, b)**. While monomeric PvPTRAMP was sufficient for binding, it showed approximately 10-fold weaker affinity (K_D_ = 292.5 ± 25.6 nM) **(Fig. 3b)**. No interaction was detected between PvRipr and PvCSS **(Fig. 3c)**, and none of the PvPCR components bound to PvCyRPA at the tested concentrations **(Fig. 3d)**. Mass photometry analysis confirmed the formation of a stable PvPCR complex, with PvPC and PvRipr each displaying monodisperse peaks when analyzed individually **(Extended data Fig. 3c)**. Formation of the PvPCR complex, following incubation of PvPC and PvRipr, was evidenced by the emergence of a higher molecular weight peak corresponding to a mass of 207 ± 29 kDa which is consistent with a 1:1:1 complex of PTRAMP:CSS:Ripr **(Fig. 3e)**. The formation of a stable complex was further validated by the co-elution of PvPC and PvRipr in size-exclusion chromatography **(Extended data Fig. 3d)**.

**Fig. 3.**
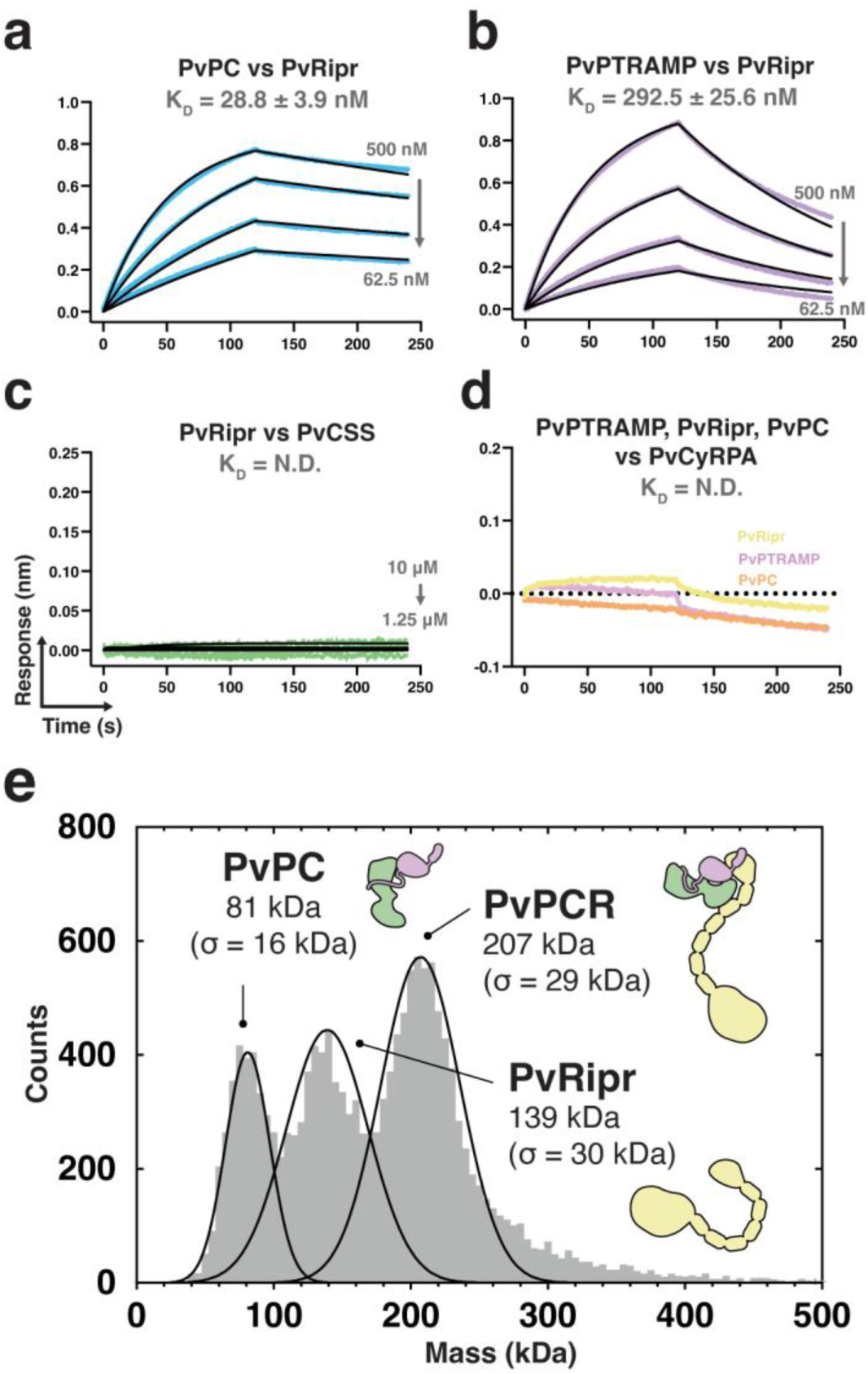
The PvPC heterodimer forms a stable complex with PvRipr. **a.**, **b.** and **c.** Representative biolayer interferometry sensorgrams of PvPC, PvPTRAMP, PvCSS, and PvRipr binding assays. Dilution series data are shown in color and 1:1 model best fit is shown in black. Representative sensorgrams of PvPTRAMP, PvRipr and PvPC binding to PvCyRPA at an analyte concentration of 5 μM. Data could not be reliably fit with a model, and so no best fit has been shown. **e.** Mass distribution of PvPC and PvRipr after pre-incubation as measured by mass photometry. Histogram data are shown in grey and Gaussian curve fit in black.

### The C-terminus of Ripr is sufficient for PTRAMP-CSS binding in multiple species of *Plasmodium*

Previous studies have demonstrated that a three-membered PTRAMP-CSS-Ripr complex is involved in *P. knowlesi* invasion^16^. We confirmed that PkPTRAMP and PkCSS form a disulfide-linked heterodimer analogous to those observed in *P. falciparum* and *P. vivax* **(Fig. 4a)** and found that the PkPC heterodimer exhibited high-affinity binding to PkRipr (K_D_ = 0.6 ± 0.1 nM) **(Fig. 4b, Extended data Fig. 4a, b).** Monomeric PkPTRAMP, but not monomeric PkCSS, was sufficient for this interaction **(Fig. 4c, d)**. Like its *P. vivax* orthologs, PkPCR formed a stable complex as demonstrated by mass photometry, with a molecular weight of 206 ± 13 kDa consistent with a 1:1:1 stoichiometry **(Extended data 4, Fig. 4e)**.

**Fig. 4.**
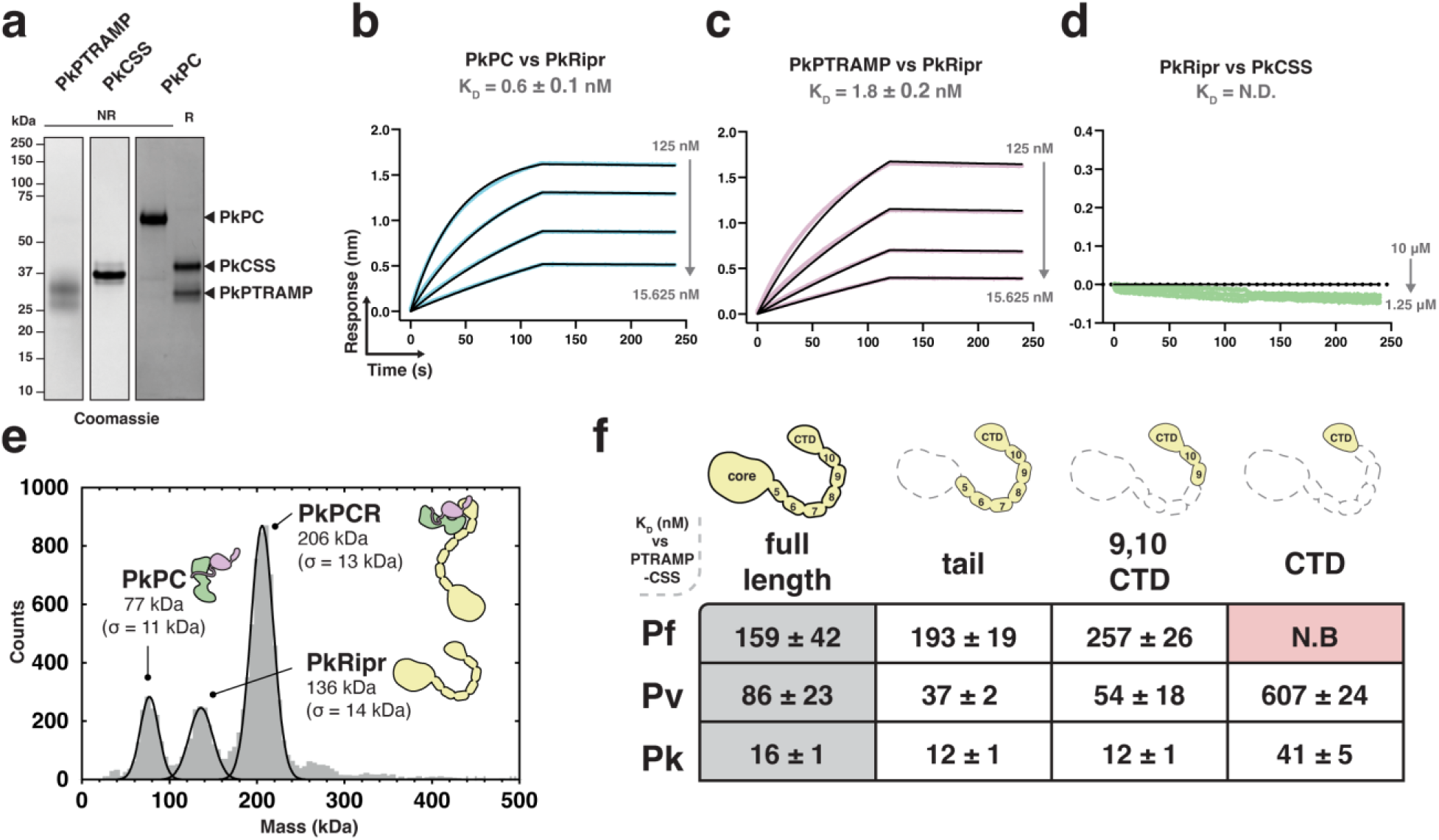
A small region of Ripr is sufficient for PTRAMP-CSS binding in *P. falciparum, P. vivax,* and *P. knowlesi*. **a.** SDS-PAGE of recombinantly expressed PkPTRAMP, PkCSS and PkPC heterodimer **b**-**d**. Representative biolayer interferometry sensorgrams of PkPC, PkPTRAMP, PkCSS and PkRipr binding assays. Dilution series data are shown in color and 1:1 model best fit is shown in black. **e.** Mass distribution of PkPC and PkRipr after pre-incubation as measured by mass photometry. Histogram data are shown in grey and Gaussian curve fit in black. **f.** The ability of Ripr and Ripr truncations to bind to PTRAMP-CSS in *P. falciparum, P. vivax* and *P. knowlesi*. The table shows the truncations used and their dissociation constant (K_D_, in nM, with standard error of the mean (SEM)) for binding their cognate heterodimer. N.B indicates no binding.

We performed a comparative biophysical analysis of PC-Ripr binding to identify the minimal regions of Ripr required for complex formation. Several truncations in the Ripr tail region were generated for *P. falciparum*, *P. vivax*, and *P. knowlesi* proteins and assessed for their ability to bind their cognate PC heterodimer **(Extended data Fig. 5)**^19^. The tail region of Ripr, which encompasses epidermal growth factor (EGF)-like domains 5-10 and the C-terminal domain (CTD), was sufficient for heterodimer binding in all three species with no observable impact on affinity **(Fig. 4f, Extended data Fig. 5)**^19^. A shorter construct containing only EGFs 9 and 10 plus the CTD also retained the ability to bind the heterodimer **(Fig. 4f)**. Interestingly, complete removal of all EGF domains, leaving only the CTD, abolished binding for *P. falciparum* proteins but not for *P. vivax* or *P. knowlesi* **(Fig. 4f)**. These results demonstrate that heterodimer binding requires only a discrete region of Ripr, with some species-specific differences in the minimal binding requirements.

### Anti-PCR antibodies are cross-reactive and differentially inhibit *Plasmodium* spp. invasion

Plasma samples from *P. falciparum*^37^, *P. vivax*^38^ and *P. knowlesi*^39,40^ infected individuals from Thailand (Tha Song Yang) and Malaysia (Sabah) were assessed to determine the extent of patient antibody response to the components of the PCR invasion complexes **(Extended data Fig. 6)**. IgG antibodies were assessed one week after clinical presentation and compared with malaria-naïve negative controls (IgG temporal kinetics from clinical presentation, one week, and one month post infection are shown in **Extended data Fig. 6**). Significant IgG antibody reactivity was detected for *P. falciparum* patients against the PCRCR complex components with the exception of PfCyRPA **(Fig. 5a)**. Antibodies from *P. vivax* patients showed reactivity to the PvPCR components PvPC, PvCSS and PvRipr, but not PvPTRAMP **(Fig. 5a)**. Similarly, antibodies from *P. knowlesi* patients showed reactivity to PkPC, PkCSS and PkRipr but not PkPTRAMP. Overall, there was a consistently low response to monomeric PTRAMP compared to other antigens and a consistently high response to monomeric CSS suggesting that the antibody response to the PC heterodimer is predominantly against CSS. High antibody responses were also observed for Ripr from all species and is consistent with Ripr being immuno-dominant as reported previously^41^. Furthermore, antibody responses to CSS and Ripr were broadly cross-reactive, with the antigens of all three species cross-reacting with sera from individuals independent of infective species **(Extended data Fig. 6)**.

**Fig. 5.**
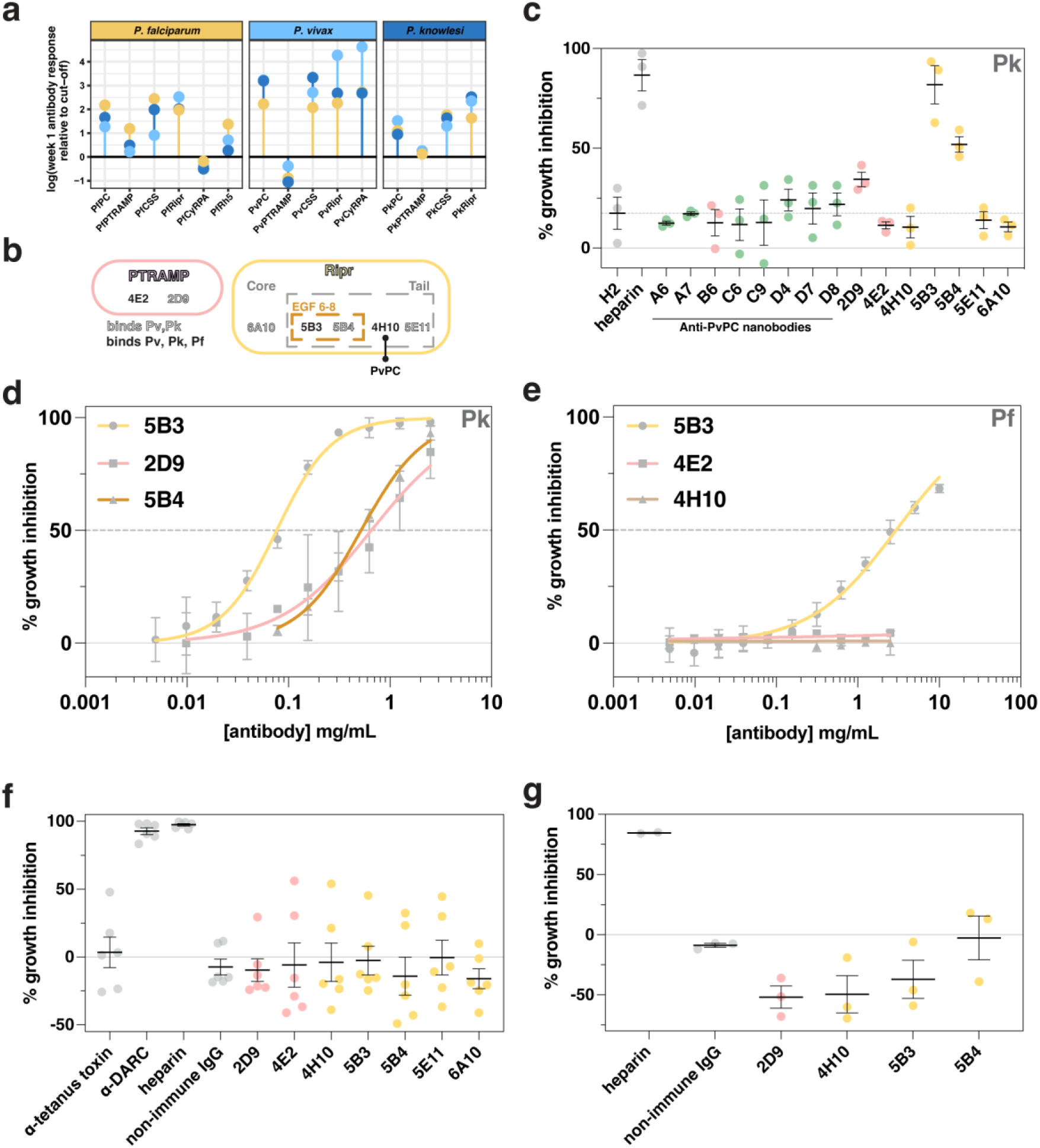
Cross-reactive antibodies targeting PTRAMP, CSS, and Ripr exhibit differential inhibition in multiple species of *Plasmodium*. **a.** IgG antibodies in human patients with *Plasmodium* infections. Fold-change peak week one antibody response relative to the seropositivity cut-off (mean of negative controls + 2x standard deviation). The panels are facetted by species of the recombinant protein (*P. falciparum*, *P. knowlesi* and *P. vivax*). The colored dots represent the plasma samples in which the proteins were assayed. **b**. Purified mouse monoclonal antibodies mapped by binding region. Open text represents antibodies that are able to bind both *P. vivax* and *P. knowlesi,* and closed text represents cross-reactivity between *P. vivax, P. knowlesi* and *P. falciparum*. Bold line indicates that 4H10 blocks PvPC binding to Ripr **c.** *P. knowlesi* growth inhibition assay of anti-PvPC and anti-PvRipr biologics. The non-inhibitory and PfCSS nanobody H2 was included as a negative control^18^. Three independent experiments were performed, and the mean and SEM are shown in black. Antibodies and nanobodies were tested at a final concentration of 0.5 mg/mL. Data points are colored according to antigen, with CSS in green, PTRAMP in pink and Ripr in yellow **d.** Growth inhibition dilution series for inhibitory antibodies in *P. knowlesi*. Growth inhibition (%) is the mean of four independent experiments. Error bars represent standard deviation. **e.** Growth inhibition dilution series for inhibitory antibodies in *P. falciparum*. Growth inhibition (%) is the mean of four independent experiments for 5B3, and two independent experiments for 4E2 and 4H10. Error bars represent standard deviation. **f.** *Ex vivo* growth inhibition assay of *P. vivax* parasites. Antibodies were tested at a final concentration of 0.5 mg/mL. Anti-Duffy antigen receptor for chemokines (DARC) mouse monoclonal antibody 2C3 was used as a positive control^46^. Data are from six independent experiments. Error bars show mean and SEM. Data points colored as in c. **g.** *P. cynomolgi* growth inhibition assay of anti-PvPC and anti-PvRipr antibodies. Antibodies were tested at a final concentration of 0.5 mg/mL. Three independent experiments were performed, and the mean and SEM are shown in black. Data points are colored as in c.

We sought to investigate the potential of antibodies and nanobodies to inhibit growth of multiple *Plasmodium* species, given the serological cross-reactivity observed. Monoclonal antibodies (mAbs) and nanobodies generated against PvPC and PvRipr were evaluated for cross-reactivity with their *P. falciparum* and *P. knowlesi* orthologs **(Extended data Fig. 7, 8, Supp. Fig. 5)**. All tested antibodies bound to PkPC and PkRipr, with three out of seven showing cross-reactivity across all three species **(Extended data Fig. 7, 8,** **Fig. 5b**)^42^. All anti-PvPC nanobodies were cross-reactive with PkPC however this cross-reactivity was much lower against PfPC with only one out of eight binding PfPC **(Supp. Fig. 5)**. Growth inhibition assays (GIAs) were performed against *P. knowlesi* to assess the inhibitory potential of anti-PvPC nanobodies and anti-PvPCR antibodies. Initial screening revealed that two anti-Ripr antibodies (5B3 and 5B4) and one anti-PC antibody (2D9) inhibited parasite growth at 0.5 mg/mL **(Fig. 5c)**. This inhibition was dose-dependent, with 5B3 and 5B4 showing half-maximal effective concentrations (EC_50_) of 77 µg/mL and 520 µg/mL respectively, while 2D9 exhibited an EC_50_ of 657 µg/mL **(Fig. 5d)**. The cross-reactive mAb 5B3 also inhibited *P. falciparum* growth, albeit with a significantly higher EC_50_ of 3 mg/mL **(Fig. 5e)**. This reduced inhibitory effect may be attributed to varying affinities of 5B3 for different Ripr orthologs **(Extended data Fig. 8e)**. Neither 4E2 nor 4H10 affected *P. falciparum* parasite growth, consistent with their lack of inhibitory activity against *P. knowlesi*. The mAb 4H10, which binds to the Ripr tail region and competes for PvPC binding **(Fig. 5b, Extended data Fig. 8c)**, showed no inhibitory activity. This suggests that the PCR complex forms prior to merozoite surface exposure, as observed previously for the PCRCR complex^18^.

Following screening of the antibodies in *P. knowlesi* and *P. falciparum* GIAs **(Fig. 5c-e, Supp. Fig. 6)**, we assessed their potential inhibitory effect on *P. vivax* merozoite invasion and parasite growth in *ex vivo* GIAs. Assays performed on Cambodian *P. vivax* parasites revealed no inhibitory effect for any of the tested antibodies **(Fig. 5f)**. To validate the *P. vivax* results, we evaluated a subset of these antibodies for their ability to inhibit growth in the closely related species *P. cynomolgi*^43^. The data from these assays were consistent with the *P. vivax* findings, confirming that none of the antibodies could inhibit parasite growth in either of these two species **(Fig. 5g)**. These results demonstrate that while antibodies against the PCR complex may exhibit cross-reactivity across recombinant PCR complexes from multiple species of *Plasmodium*, this cross-reactivity does not necessarily correlate with growth inhibitory capacity. As these antibodies were raised against the *P. vivax* protein, these results either suggest minor functional differences between the complexes of *P. falciparum* and *P. knowlesi* compared to *P. vivax* and *P. cynomolgi* or that the PCR components are less critical for invasion of *P. vivax* and *P. cynomolgi*.

### Cryo-EM analysis of PkPCR supports AlphaFold predictions

To provide more confidence in the predicted models of the PCR complexes, and to understand how inhibitory antibodies may function, we carried out cryo-electron microscopy (cryo-EM) experiments on the PkPCR complex. Cryo-EM analysis of the PkPCR^tail^ complex revealed an overall shape consistent with the AlphaFold prediction **(Fig. 6a, Extended data Fig. 9)**. The addition of the antigen-binding fragment (Fab) of 5B3 allowed unambiguous assignment of the orientation of the two-dimensional (2D) classes **(Fig. 6a)**. Furthermore, comparison of Fab bound and unbound classes showed no discernible differences in the PCR complex which confirmed that 5B3 binds to the tail region of Ripr without interfering with complex formation **(Fig. 5b, Extended data Fig. 8, 9)**. This suggests that parasite inhibition by 5B3 likely has a direct effect on Ripr function during invasion rather than on the complex as a whole. The cryo-EM data, combined with the PvPC crystal structure, strongly support the predicted PCR complex structure **(Fig. 6b)**. In this model, the PTRAMP-CSS heterodimer is formed by an intermolecular disulfide bond. This heterodimer engages Ripr via two interfaces: PTRAMP clinching the CTD of Ripr, and the D2 domain of CSS interacting with EGF 9 of Ripr **(Fig. 6b)**. The remaining mass of Ripr likely extends below the PCR complex, where it may interact with other invasion proteins (such as PfCyRPA) or potentially with erythrocyte proteins.

**Fig. 6.**
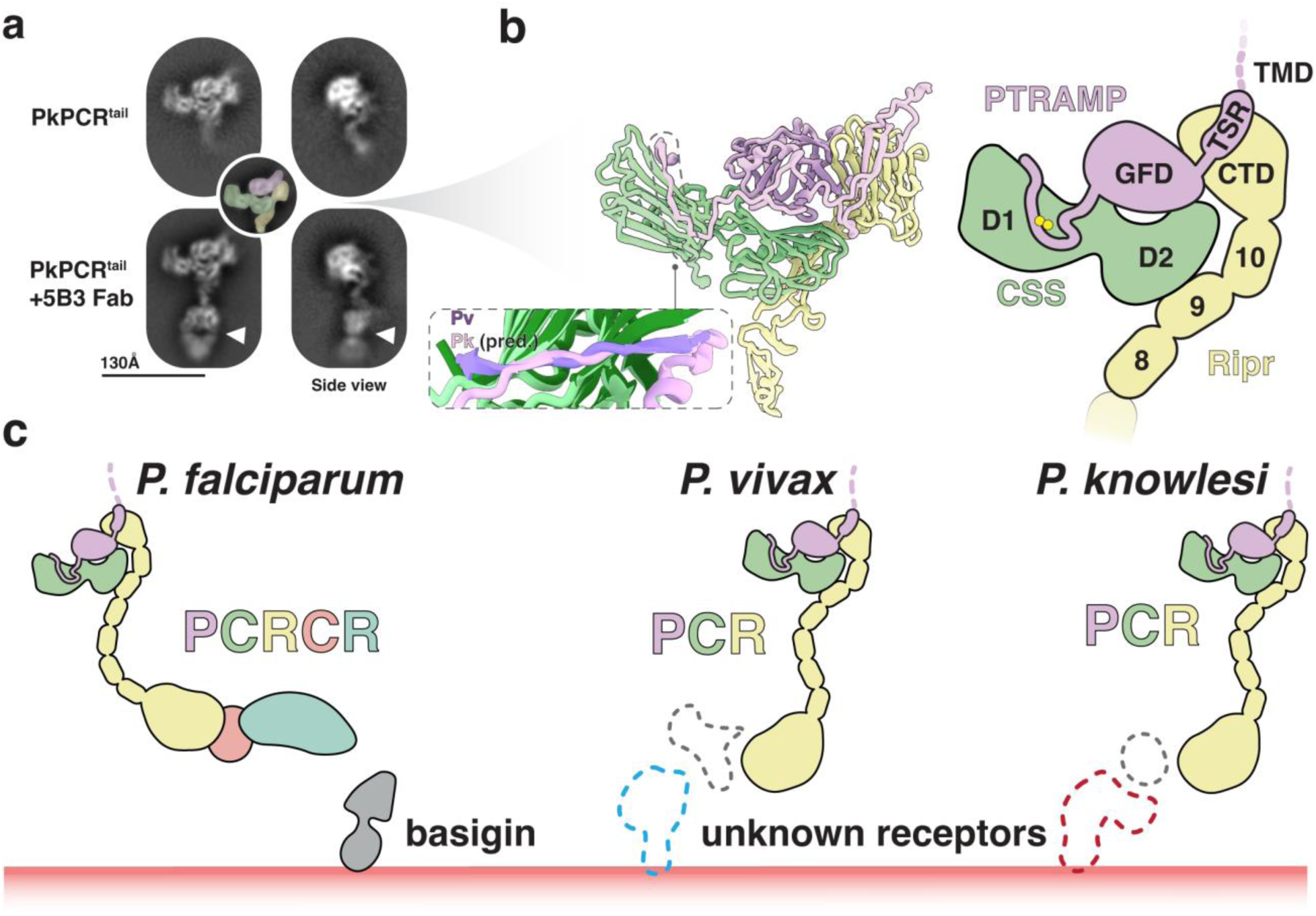
PTRAMP, CSS, and Ripr form a core invasion scaffold in *Plasmodium* spp. **a.** Cryo-EM 2D class averages of the PkPCR^tail^ complex. Addition of the 5B3 Fab fragment shows distinct density at the end of the Ripr tail (white triangle). Inset shows the PkPCR^tail^ complex colored according to the predicted model **b.** AlphaFold 3 predicted model (left) and diagram (right) of PkPCR^tail^. Transmembrane domains, signals sequences and large disordered regions have been omitted for clarity. Inset shows the alignment of the PvPC (purple and dark green) structure and the PkPC predicted structure (pink and light green) **c.** Model of the PCRCR complex of *P. falciparum* and PCR complexes of *P. vivax,* and *P. knowlesi.* PTRAMP, CSS and Ripr form a conserved three-membered complex that serves as a scaffold for an invasion complex. In *P. falciparum* this complex involves CyRPA and Rh5. In *P. vivax* and *P. knowlesi* this complex likely contains other components (dashed grey) that have yet to be identified, which engage with the erythrocyte membrane via host-cell specific receptors.

While PfPC lacks the ability to bind erythrocytes directly, it enhances Rh5 binding when incorporated into the PCRCR complex^18^. Previous studies have shown that PkPTRAMP can bind erythrocytes; however, these experiments were performed with monomeric PkPTRAMP and not heterodimeric PkPC^16^. We performed flow-cytometry based erythrocyte binding assays to assess whether PkPC or the PkPCR complex bound to erythrocytes. Neither PkPC nor PkPCR showed significant binding to erythrocytes relative to the positive control, PfRh5 **(Extended data Fig. 10a, b)**. Considering that *P. vivax* invades reticulocytes exclusively, we extended our investigation to include reticulocyte binding assays. Similarly, we observed no binding of PvPC or PkPC to reticulocytes **(Extended data Fig. 10c, d)**. Collectively our analysis found no evidence of erythrocyte or reticulocyte binding by PkPC, PvPC, or the PkPCR complex.

We therefore hypothesize that PTRAMP, CSS and Ripr form a core invasion scaffold in *Plasmodium* parasites. This scaffold provides the basis for the assembly of species-specific complexes that are adapted for binding a diverse set of host erythrocyte receptors **(Fig. 6c)**. In *P. falciparum* this complex incorporates CyRPA and Rh5 which facilitate invasion via the essential interaction with basigin. The equivalent proteins in *P. vivax* and *P. knowlesi* that are responsible for erythrocyte binding are yet to be identified. If findings from *P. falciparum* are applicable to these species, these interactions will be an essential step in merozoite invasion.

## Discussion

The highly conserved nature of PTRAMP, CSS, and Ripr across the *Plasmodium* genus, combined with their demonstrated essential roles in both *P. falciparum*^18^ and *P. knowlesi*^16^ invasion, positions these proteins as compelling targets for understanding fundamental mechanisms of merozoite invasion. Recent advances in protein structure prediction through AlphaFold^33^ have enabled a comprehensive comparative analysis of these proteins across three clinically significant *Plasmodium* species: *P. falciparum*, *P. knowlesi*, and *P. vivax*. Our cross-species structural and functional analyses reveal that PTRAMP, CSS, and Ripr form a conserved invasion scaffold in *Plasmodium* parasites that serves as a foundation for the assembly of species-specific protein complexes **(Fig. 6c)**. These complexes appear to be evolutionarily adapted for engaging diverse host erythrocyte receptors. In *P. falciparum*, this complex includes CyRPA and Rh5, which mediate the essential interaction with the host receptor basigin. While the equivalent erythrocyte-binding proteins in *P. vivax* and *P. knowlesi* remain unidentified, the conservation of this core scaffold suggests that analogous essential receptor-ligand interactions likely govern invasion in these species.

Structural analysis of PvPC revealed a critical intermolecular disulfide bond between PvPTRAMP and PvCSS. The essentiality of this linkage was previously established in *P. falciparum* invasion^18^, and the evolutionary conservation of these cysteine residues across *Plasmodium* species strongly suggests that PTRAMP-CSS heterodimerization represents a fundamental feature throughout the genus. Optimization of recombinant PfPC heterodimer revealed a much tighter interaction with PfRipr than previously reported^18,19^, aligning with the structural architecture predicted by AlphaFold ^33^. Our biochemical studies demonstrated that the heterodimeric PkPC forms a high-affinity complex with PkRipr, corroborating earlier pull-down mass spectrometry data from *P. knowlesi* parasites^16^ and providing robust evidence for the biological significance of this complex *in vivo*. The formation of this trimeric complex extends beyond *P. knowlesi* and *P. falciparum*, as we also demonstrated its assembly in *P. vivax*, providing evidence for a conserved molecular feature across multiple *Plasmodium* species.

Previous studies have shown that while PfRipr’s core region interacts with CyRPA to form the RCR complex, its C-terminal tail mediates PfPC binding^17,19^. Our findingshave further refined this understanding by demonstrating that only a small domain within the Ripr tail is required for PC heterodimer binding. In *P. falciparum*, this binding region encompasses EGFs 9 and 10 and the CTD of PfRipr. Notably, in both *P. vivax* and *P. knowlesi*, the CTD alone is sufficient for PC heterodimer binding, indicating evolutionary divergence in these interactions across *Plasmodium* species. The observation that both PvPC and PkPC can bind Ripr^CTD^, coupled with the finding that PvPTRAMP and PkPTRAMP alone are sufficient for Ripr binding, suggests that the PTRAMP- Ripr^CTD^ interaction serves as the primary interface driving complex formation.

Analysis of antibodies from patients infected with *P. falciparum*, *P. knowlesi*, or *P. vivax* has revealed significant cross-reactivity of antibodies to CSS and Ripr across these *Plasmodium* species. It is unlikely that these patients had previously been recently infected with all three *Plasmodium* species, particularly given the low transmission in these settings, suggesting that antibodies generated against the PCR complex targeted conserved epitopes. This raised the potential for cross-species antibody-mediated inhibition of invasion. Indeed, analysis of monoclonal antibodies identified the Ripr-binding mAb 5B3, which exhibited cross-inhibitory activity against both *P. knowlesi* and *P. falciparum*, but not *P. vivax* or *P. cynomolgi*. Interestingly, this mAb was raised against PvRipr, suggesting functional differences with the PCR complex between these *Plasmodium* species that may render the conserved epitope on PvRipr and PcRipr inaccessible to the antibody in the full complex. It is also possible that the essentiality of the complex differs between species. The high degree of conservation of the complex and its components across the *Plasmodium* genus would make this conclusion unlikely. The differential inhibition is unlikely to be due to antibody affinity but may reflect differences in how Ripr functions during invasion of different host cells. The identification of cross-species neutralizing antibodies is an attractive finding for vaccinology^44,45^; however, the inhibitory activity of such naturally acquired cross-reactive antibodies is yet to be fully explored. The conservation of the PCR complex makes it an attractive target for such an approach.

The considerable length of Ripr (>150 Å), while largely uninvolved in PCR complex formation, may be important for enabling the PCR/PCRCR complex to bridge the gap between the merozoite surface and host cell membrane during invasion **(Fig 6c)**^19^. This model is supported by previous studies showing that antibodies targeting EGF domains 6, 7, and 8 within the Ripr tail effectively inhibit parasite growth *in vitro*^24,41^. Our findings suggest these antibodies may function by preventing adequate extension of Ripr between the two membranes, thereby disrupting receptor engagement. Understanding the structural basis of Ripr inhibition will be crucial for elucidating its role within the PCR complex and its potential as a therapeutic target.

## Acknowledgements

The authors thank Australian Red Cross Blood Service and Bone Marrow Donor Institute (BMDI) Cord Blood Bank for blood. We acknowledge Professor Jamie Rossjohn and the Monash Macromolecular Crystallisation Facility (https://www.monash.edu/researchinfrastructure/mmcp), where crystallization screening was undertaken. This research was undertaken in part using the MX2 beamline at the Australian Synchrotron, part of the Australian Nuclear Science and Technology Organisation, and made use of the Australian Cancer Research Foundation (ACRF) detector. We thank Professor Wai-Hong Tham for supply of the PvRBP2b plasmid and anti-PvRBP2b sera. We acknowledge all field teams in Thailand and Malaysia who contributed to collection of the used samples. We acknowledge the VBDR at WEHI for collection of Melbourne controls. This work was supported by the Gates Foundation (INV-074041), National Health and Medical Research Council of Australia (NHMRC) (grants 637406, APP1173049, GNT1173210), Drakensburg Trust, Australian Research Council (ARC FT240100420 University of Adelaide Research Scholarship), National Institutes of Health (NIH 5R01AI140751), and Victorian State Government Operational Infrastructure Support grant.

## Author Contributions

BAS designed experiments, expressed proteins, performed and analyzed biophysical experiments, solved the crystal structure with assistance from SWS, analyzed cryo-EM data, and wrote the manuscript. PSL, LBFD, KHL and SD performed parasite growth inhibition assays and analyzed data. XX and NCJ purified proteins and performed biophysical assays. AA and PSL performed serological assays and analyzed data. TW, MJG, NMA, JS and RJL organized and collected patient plasma and clinical data for antibody analysis. AL carried out cryo-EM data collection. RJL, MJG, MTD, JP, DWW, SWS designed and interpreted experiments. AFC and SWS designed and interpreted experiments and wrote the manuscript. All authors read and edited the manuscript.

## Materials and Methods

### Recombinant protein expression

All gene sequences used were retrieved from the VEuPathDB (accessed through www.plasmodb.org)^47^ from reference strains (3D7 for *P. falciparum,* PvP01 for *P. vivax (*with the exception of PvRBP2b for which the *Sal-*1 sequence was used), and strain H for *P. knowlesi*). All genes were synthesized by Genscript (Singapore) unless otherwise stated.

#### P. falciparum

Recombinant PfRh5, PfCyRPA, PfPTRAMP, PfPC and PfRipr constructs were produced as described previously, with some changes made to the synthetic gene constructs used for expression^18^. PfPTRAMP, comprising residues 31 to 307, was subcloned into the pAcGP67a vector with a C-terminal C-tag. Four potential N-linked glycosylation sites were removed, at positions Asn112, Asn149 and Asn155 by mutation to Gln, and at position Asn195 by mutation of Thr197 to Ala, to produce PfPTRAMP_31-307_4x. To produce a large amount of pure monomeric PfPTRAMP, another construct was made that contains all of the same mutations and also contains Cys60Ser mutation to prevent disulfide formation, termed PfPTRAMP_31-307_4xC60S. PfCSS was subcloned into the pAcGP67a vector with a C-terminal FLAG-tag preceded by a TEV protease cleavage site. This construct has all six potential N-linked glycosylation sites removed at positions Asn74, Asn88, Asn192, Asn234, Asn261 and Asn283 by mutation of Ser76, Thr90, Ser194, Thr236 and Thr263 to Ala and Asn283 to Gln, to produce PfCSS_21-290_6x^18^. The previous PfRipr construct^18^ was altered with the following mutations: Thr966Ala-Ser1023Ala yielding the construct PfRipr_20-1086_2xA. PfRipr^tail(^aa 717-1086), PfRipr^EGF 9,10,CTD^(aa 899-1086) and PfRipr^CTD^(aa 981-1086), were all synthesized by Genscript and purified in an identical fashion to PfRipr.

#### P. vivax

The *pvptramp* gene (PVP01_1436800, aa 21-297) excluding the transmembrane and cytoplasmic domains was subcloned into a modified pTRIEX2 vector that contains an N-terminal Small Ubiquitin-like Modifier (<u>S</u>UMO)-<u>F</u>lag tag followed by a Tobacco Etch Virus (<u>T</u>EV) protease cleavage site (from here on termed SFT). Potential N-glycosylation sites were assessed and one site, Asn115, was removed by mutation of Ser117 to Ala. This yielded SFT_PvPTRAMP_21- 297_S117A. This construct was then further cloned to incorporate a C-terminal Avitag, yielding SFT_PvPTRAMP_21-297_S117A_Avi. Both constructs were expressed in Human Embryonic Kidney (HEK) Expi293F cells (Life Technologies) as secreted soluble proteins. Transient transfection was carried out as per the manufacturer’s protocol and the culture medium harvested 5-6 days post-transfection. The proteins were purified via multiple rounds of binding and eluting using Anti-Flag M2 Affinity Gel (Merck) and 100 μg/mL of Flag peptide (Genscript) in HBS (20 mM HEPES pH 7.2, 150 mM NaCl). The eluted fractions were pooled and incubated with TEV protease (1 mg of TEV for every 10 mg of protein) overnight at 4°C. His-tagged TEV was removed by applying the protein solution to nickel-nitrilotriacetic acid (Ni-NTA) agarose resin (Qiagen) and collecting the flowthrough. The flowthrough was then concentrated on a 10,000 dalton (Da) Molecular Weight Cut-Off (MWCO) Amicon Ultra-15 Centrifugal Filter (Merck) and applied to an S75 Increase 10/300 column (Cytiva) connected to an Akta Pure (Cytiva) to separate the TEV-cleaved PvPTRAMP from the SUMO-Flag tag. Peak fractions were assessed for purity via sodium dodecyl sulfate–polyacrylamide gel electrophoresis (SDS-PAGE) and pure fractions were pooled and concentrated. Glycerol was added (10% v/v final) to the concentrated protein and then aliquoted and flash frozen in liquid nitrogen (LN_2_).

The full-length *pvcss* gene (PVP01_1344100, aa 22-381) was subcloned into a modified pTRIEX2 vector with a C-terminal <u>F</u>lag-tag preceded by a <u>T</u>EV protease cleavage site (from here on termed TF). N-glycosylation sites were predicted and three potential sites, Asn114, Asn180, Asn352, were mutated via three mutations: Ser116Ala, Thr182Ala and Ser354Ala. This yielded PvCSS_22- 381_S116A_T182A_S354A_TF. This construct was further cloned to remove a predicted N-terminal repeat region to aid in crystallisation. This yielded the construct PvCSS_115- 381_S116A_T182A_S354A_TF. These constructs were expressed as soluble secreted proteins in HEK Expi293F cells (Life Technologies) as above.

To generate disulfide-linked PvPTRAMP-PvCSS (PvPC), PvPTRAMP and PvCSS were co-expressed in HEK Epi293F cells (Life Technologies) at a ratio of 50:50 PvPTRAMP:PvCSS for full length CSS and a ratio of 40:60 PvPTRAMP:PvCSS for PvCSS_115. The expression and purification were carried out as above with size-exclusion chromatography performed using either an S200 Increase 10/300 GL (Cytiva) or S200 16/600 HiLoad (Cytiva).

The full-length *pvripr* gene (PVP01_0816800, aa 22-1074) was subcloned into pAcGP67a with a C-terminal 6xHis-tag yielding PvRipr_22-1074_His and expressed in Sf21 cells using the flashBAC ULTRA baculovirus system (Oxford Expression Technologies). After initial transfection, the P1 virus was amplified and titrated several times to produce a P3 virus. This P3 virus was used for large scale expression. The proteins were expressed as soluble secreted proteins. The culture medium was harvested 3 days after the addition of P3 virus. The media was concentrated via tangential-flow filtration with a 3,000 Da MWCO (Merck) to reduce the volume ∼10-15 fold. This resultant concentrate was then dialysed into TBS (20 mM Tris pH 8.5, 150 mM NaCl) overnight at 4°C. Imidazole was then added to a final concentration of 10 mM and the culture media passed over Ni-NTA Agarose resin (Qiagen), washed with TBS + 20 mM imidazole, and eluted in TBS + 500 mM imidazole. The eluted protein was then concentrated on a 30,000 Da MWCO Amicon Ultra-15 Centrifugal Filter (Merck) and applied to an S200 Increase 10/300 GL column (Cytiva) connected to an Akta Pure (Cytiva). Peak fractions were pooled, concentrated and flash frozen as above. PvRipr truncations (PvRipr^tail^(aa 665-1074), PvRipr^EGF 9,10,CTD^(aa 844- 1074), PvRipr^CTD^(aa 972-1074)) were synthesized via the Genscript mutagenesis service. The truncated constructs were subcloned into the pAcGP67a vector and purified in an identical manner to the full-length construct.

PvRipr^EGF 6–8^(aa 717-843) was synthesized and subcloned into pET28a yielding a construct with an N-terminal 6x His tag followed by a TEV site. Expression was carried out in *Escherichia coli (E. coli)* strain SHuffle® T7 (New England Biolabs) grown in terrific Broth with 40 μg/mL of kanamycin. One litre of culture was grown in incubators at 37°C and shaking at 180 revolutions per minute (rpm) until an optical density at 600 nm (OD 600) of around 1.0 was reached. Isopropyl ß-D-1-thiogalactopyranoside (IPTG)(Astral) was then added to a final concentration of 1mM, and protein expression was continued at 16°C for 16-18 hours. Cells were then harvested via centrifugation and the pellet resuspended in TBS pH 8.5, and with cOmplete ethylenediaminetetraacetic acid (EDTA)-free protease inhibitor cocktail (Roche). The resuspended cells were then sonicated, and the cellular extract clarified by centrifugation at 30,000 x*g* for 30 minutes at 4°C. Imidazole was added to the clarified supernatant to a final concentration of 10 mM and then passed over pre-equilibrated Ni-NTA resin, washed with TBS pH 8.5 + 20 mM imidazole, and then eluted in TBS pH 8.5 containing 500mM imidazole. The eluted protein was then concentrated on a 10,000 MWCO Amicon Ultra-15 Centrifugal Filter (Merck) and injected onto an S200 16/600 HiLoad (Cytiva) equilibrated in TBS pH 8.5. Peak fractions were then concentrated, supplemented with glycerol to a final concentration of 10% (v/v) and flash frozen in liquid nitrogen.

The *pvcyrpa* gene (PVP01_0532400, aa 24-362) was subcloned into pTRIEX2-TF vector which yielded PvCyRPA_22-362_TF. Two predicted N-glycoslyation sites, Asn78 and Asn282, were removed with the following mutations: Thr80Ala, Thr284Ala. The protein was expressed in HEK Expi293F cells and purified in an identical manner to PvPTRAMP above.

PvRBP2b (PVX_094255, aa 161-1454) was purified as described previously^48^.

#### P. knowlesi

The *pkptramp* gene (PKNH_1437600, aa 21-297) excluding the transmembrane and cytoplasmic domains was subcloned into a modified pTRIEX2-SFT as was done for PvPTRAMP. Potential N-glycosylation sites were assessed and two sites, Asn115 and Asn261, were removed by mutation of Ser117 and Ser263 to Ala. This yielded SFT_PkPTRAMP_21-297_S117A_S263A. A PkPTRAMP construct expressing a C-terminal Avitag for biotinylation was made using PCR and restriction digests to yield SFT_PkPTRAMP_21-297_S117A_S263A-Avi. Expression of PkPTRAMP and PkPTRAMP-Avi was carried out in an identical manner to PvPTRAMP described above.

The *pkcss* gene (PKNH_1353400, aa 22-362) was subcloned into pTriEX2-TF. Five potential N-glycosylation sites, Asn96, Asn161, Asn175, Asn243 and Asn333 were removed via five mutations: Ser98Ala, Thr163Ala, S177Ala, S245A and S335A. This yielded PkCSS_22-362_ S98A_T163A_S177A_S245A_S335A _TF. Expression of PkCSS was carried out in an identical manner to PvCSS above.

Expression of the PkPC heterodimer was carried out in an identical manner as described for PvPC. The ratio of PkPTRAMP:PkCSS deoxyribonucleic acid (DNA) used was 60:40 when PkPTRAMP-SFT and PkCSS-TF were being used, and 50:50 when PkPTRAMP-Avi was being used.

The *pkripr* gene (PKNH_0817000, aa 22-1096) was subcloned into pAcGP67a with a C-terminal 6xHis-tag yielding PkRipr_22-1096_His, as per the PvRipr construct. Truncations of the full-length construct were made by Genscript using the mutagenesis service, to produce PkRipr^tail(^aa 669-1096), PkRipr^EGF 9,10,CTD^(aa 848-1096) and PkRipr^CTD^(aa 994-1096). All PkRipr constructs were expressed and purified in an identical manner to the equivalent PvRipr constructs.

For all constructs containing an Avitag, *in vitro* biotinylation was carried out as previously described^49^.

### Antibodies and nanobodies

One alpaca was subcutaneously immunized six times 14 days apart with 130 μg (800 μg total) of recombinant PvPC. GERBU FAMA (GERBU Biotechnik GmbH, Heidelberg, Germany) was used as an adjuvant. Whole blood was collected three days after the last immunization for the preparation of lymphocytes. Nanobody library construction was carried out according to established methods^50^. Briefly, alpaca lymphocyte mRNA was extracted and amplified by reverse transcription PCR (RT-PCR) with nanobody-encoding, gene-specific primers. This produced a library of nanobody cDNA sequences that contained approximately 10^8^ sequences. The sequences that were cloned into the pMES4 phagemid vector were amplified in *E. coli* TG1 strain and subsequently infected with M13KO7 helper phage for downstream recombinant phage expression. Handling of the alpaca for scientific purposes was approved by Agriculture Victoria, Wildlife and Small Institutions Animal Ethics Committee, project approval No. 26-17.

Biopanning was performed over two rounds with 1 μg of immobilized antigen as previously described^50^. Ninety-four positive clones were taken for further screening via enzyme-linked immunosorbent assay (ELISA). Clones showing positive binding by ELISA (n = 93) were sequenced. Of these, 71% were full length Variable Heavy domain of Heavy chain (VHH) (n = 66).

Nanobodies were expressed in the periplasm of *E. coli* WK6 cells as described previously (6). Briefly, bacteria (250 mL) were grown in Terrific Broth at 37°C to an OD 600 of 0.7. The cultures were then induced with 1 mM IPTG (Astral) and grown overnight at 28°C. Cells were harvested and resuspended in PBS containing 20% sucrose and 20mM imidazole to rupture the periplasm. EDTA was added to a final concentration of 5 mM, and the cells were incubated on ice. MgCl_2_ was then added to a final concentration of 10 mM, and the periplasmic extract was harvested via centrifugation. The nanobodies were purified via standard Ni-NTA purification methods.

Monoclonal antibodies were raised in mice as per the Walter and Eliza Hall Animal Ethics Committee approved procedures. All monoclonal antibodies were produced by the WEHI Antibody Facility. Mice were injected with 80-180 μg of protein three times and then boosted once with 30-60 μg. After cloning of hybridomas, the supernatants were tested via ELISA and BLI. Based on these results, several hybridomas for each antigen were selected for further scale up of purified immunoglobulin G (IgG).

### Structure prediction

Prediction of PCR complexes from multiple *Plasmodium* species was done using the AlphaFold 3 server^34^.

### Biolayer interferometry (BLI)

Biolayer interferometry (BLI) experiments were carried out on an Octet Red96e (Sartorius) at 25°C. For kinetics analysis ligands were immobilized onto either anti-penta-His (His1K), streptavidin (SAX or SAX2) or Ni-NTA (NTA) biosensors (Sartorius) depending on the affinity tag present on the protein (His-tag or biotinylated Avitag). Ligands were diluted to 10-40 μg/mL in 1x kinetics buffer (PBS, pH 7.4, 0.1% (w/v) bovine serum albumin (BSA), 0.02% (v/v) Tween-20) prior to immobilisation. Biosensors were initially dipped in kinetics buffer for 30-60 seconds to establish a baseline signal, and then dipped into wells containing the ligand, followed by another 30-60 second baseline. After the second baseline step, the ligands were then dipped into wells containing two-fold dilution series of analyte. Association was measured for 120 seconds and then the biosensors were dipped into kinetics buffer to measure the dissociation for another 120 seconds. Data were analysed using Sartorius Data Analysis software 11.0. Kinetic curves were fitted using a 1:1 binding model.

Competition studies for anti-PvPC nanobodies were performed using Ni-NTA (NTA) biosensors (Sartorius) with His-tagged nanobodies as the ligand (diluted to 5 μg/mL in kinetics buffer). After a 30 second baseline step, the biosensors were dipped into wells containing an irrelevant nanobody that does not bind to PvPC to quench the biosensor and ensure no free sites are present for the downstream steps. Following a second baseline step, the biosensors were dipped into PvPC diluted to 500 nM in kinetics buffer. After loading of PvPC onto the biosensors, a final baseline step was performed before the biosensors were dipped into either secondary nanobody (at 10 μg/mL diluted in kinetics buffer) or PvRipr (at 200 nM diluted in kinetics buffer). Data were analysed using Sartorius’ Data Analysis software 11.0 and the epitope bins were assessed by normalization and manual curation.

Antibody kinetics were determined similarly to the above methods. Anti-Mouse IgG Fc Capture (AMC) biosensors (Sartorius) were used to immobilize mouse monoclonal antibodies at a concentration of 5-20 μg/mL in kinetics buffer. Antibody competition studies were carried out in a similar manner to the nanobodies, however anti-pentaHis (His1K) biosensors were used.

### Protein crystallization

PvPC_115 was purified as above. PvPC and nanobody D7 were co-complexed with the nanobody at 3x molar excess. The free nanobody was separated from the PvPC-nanobody complex by size-exclusion chromatography on an S200 Increase GL 10/300 (Cytiva) in HBS. Peak fractions were pooled and concentrated to ∼5-6 mg/mL and set up in coarse screen sitting drop crystal trays at the Monash Macromolecular Crystallisation Facility. Needle-like crystals formed after ∼9 days in 0.2M ammonium sulfate ((NH_4_)_2_SO_4_) and 20% (w/v) polyethylene glycol (PEG) 3,350. Further in-house optimization of conditions yielded large crystals in 0.2M (NH_4_)_2_SO_4_ and 16%(w/v) PEG- 3,350. Crystals were looped in mother liquor containing 10% (v/v) glycerol and flash frozen in liquid nitrogen. Diffraction data were collected with the MX2 beamline at the Australian Synchrotron (Clayton, Australia) at 100 K (λ = 0.9537 Å). Statistics are in Table 1.

### Structure determination and model building

Diffraction data were processed with the XDS package^51^ before being scaled and merged using Aimless^52^ in the CCP4 suite^53^. The program Matthews^54^ was used to estimate the number of molecules in the asymmetric unit. An AlphaFold 2^33^ model of PvCSS constituting residues 115- 381 was used as a search model for molecular replacement using Phaser^55^. After 3 copies of PvCSS were fitted, additional searches were performed with a nanobody structure. To ensure the best fit possible, a BLASTp (https://blast.ncbi.nlm.nih.gov/Blast.cgi?PAGE=Proteins) search was performed with nanobody D7 to find the structure with the highest sequence similarity for molecular replacement searches. This search yielded a nanobody (PDB: 7N0R), which was used as a search model with the complementarity determining region 3 (CDR3) sequence removed ^56^. The structure was then iteratively refined in Phenix^57^ and assessed and modified with Coot^58^. Clear density extended from the unpaired cysteine in PvCSS, C122, that was not accounted for by either PvCSS or nanobody D7. Due to the crystals being set up with the PvPC heterodimer, it was reasonable to conclude that this density belonged to PvPTRAMP. PvPTRAMP residues 41-53 were built into the electron density *de novo* independently for each of the three molecules in the asymmetric unit. Refinement and model statistics are described in Table 1. For analysis of the contacts formed between PvPTRAMP and PvCSS, and PvCSS and D7, the program Contact (part of CCP4 suite)^53^ was used in conjunction with PISA server (https://www.ebi.ac.uk/pdbe/pisa/) and are summarized in Tables 2 and 3. Nanobodies were renumbered according to the Kabat numbering system, as determined by ANARCI^59^.

### Mutiple sequence alignment

Multiple sequence alignments were computed using ESPript 3.0^60^.

### Mass photometry

Mass photometry experiments were carried out on a Two^MP^ mass photometer (Refeyn). Each well was focused after the addition of 10 μL of filtered PBS. Once focused, 10 μL of protein at either 50 nM (*P. knowlesi*) or 100 nM (*P. vivax*) was added, mixed, and events were recorded for one minute using AcquireMP (Refeyn). Raw data processing was done in DiscoverMP (Refeyn) and the data exported and presented using Prism v9 (GraphPad). In-house recombinant mouse mixed lineage kinase-like (MLKL), human glutamine synthetase and human catalase were used for the construction of a calibration curve.

### Human samples

Human plasma samples were utilized from three cohorts of *Plasmodium* infected patients along with three cohorts of malaria-naïve negative controls. Patients infected with *P. vivax* were recruited from Tha Song Yang, Thailand, during the year 2014, as previously described ^38^, with a subset of 34 included in the current study. Patients infected with *P. falciparum*^37^ and *P. knowlesi*^39,40^ were recruited from Sabah, Malaysia, during the years 2010-2018 and 2012-2014, respectively. *P. falciparum* (n = 31) and *P. knowlesi* (n = 33) infected patients were included in the current study. Plasma samples were assayed from time of clinical presentation, 1 week later, and 1 month later. 28 malaria-naïve samples from the Melbourne Volunteer Biospecimen Donor Registry (VBDR) and 29 malaria-naïve samples from the Thai Red Cross (TRC) were utilized to create seropositivity cut-offs. Individuals from the TRC donated blood in Bangkok, a malaria-free region of Thailand, and had not had malaria diagnosed in the year prior nor had they travelled to endemic regions in the prior three years, as previously described ^61^. An additional set of afebrile healthy controls (n=30) were assayed from Sabah, Malaysia; however, these individuals may have had prior *Plasmodium* infections and were thus not utilized to create the seropositivity cut-off.

Ethical approval for sample use was provided by WEHI Human Research Ethics Committee (14/02), with original study approval in Thailand (Faculty of Tropical Medicine, Mahidol University, MUTM 2014-025-01 and 02) and Malaysia (Menzies School of Health Research, HREC 12-1815, 16-2544, 10-1431, 12-1807). All individuals gave informed consent and/or assent to participate in the studies.

### Multiplexed antibody assay

Recombinant *Plasmodium* proteins were coupled to unique regions of magnetic, fluorescent, Bio-Plex microbeads (Bio-Rad) following the manufacturer’s instructions and as previously described^62^. Briefly, 200 µL of microbeads were washed then activated for 20 minutes with sulfo-N-hydrosuccinimide (50 mg/mL) and N-ethyl-N-(3-dimethylaminopropyl) carbodiimide (EDC) (50 mg/mL) in monobasic sodium phosphate (pH 6.2). Following further washing, the activated microspheres were resuspended in PBS with 1-6 µg of *Plasmodium* protein. After overnight incubation, the coupled beads were washed and then stored in PBS-TBN (PBS, 0.1% (w/v) BSA, 0.02% (v/v) TWEEN-20, 0.05% (w/v) sodium azide, pH 7) at 4 °C until further use. Microbeads were always kept protected from light.

Plasma samples were diluted in PBT (1X PBS, 1% (w/v) BSA, 0.05% (v/v) Tween-20) at a dilution of 1:100. For *P. vivax* and *P. falciparum* antigens, plasma samples from hyper-immune individuals from PNG were used as a positive control. For *P. knowlesi* antigens, plasma samples from acutely infected *P. knowlesi* patients were used as the positive control. Both positive control pools were used to create a modified reference standard curve, starting at 1:50 with a 5 point 5-fold serial dilution. Diluted samples (50 µL) all controls and patients) were added to black flat-bottom 96- well plates and mixed with 50 µL of the coupled-antigen bead mixture (0.1 µL of each coupled antigen per well in PBT), then incubated for 30 minutes. The plate was washed and then 100 µL of 1:100 phycoerythrin (PE)-conjugated anti-human secondary antibody (Jackson Immunoresearch) was added and incubated for a further 15 minutes. Plates were washed then resuspended in PBT before being read on a MAGPIX instrument. Median fluorescent intensity was converted to arbitrary relative antibody units (RAU) using the standard curves, to adjust for plate-plate variation^61^.

### Statistical analysis

An antigen-specific seropositivity cut-off was set as the mean of the negative controls (VBDR + TRC) plus two times the standard deviation. Data are presented as the fold change of the mean peak week 1 antibody response relative to the seropositivity cut-off. RAU values of samples and control cohorts are shown in Extended Data.

### Growth inhibition assays

*P. knowlesi* growth inhibition assays were undertaken using *P. knowlesi* YH1 parasites over 2 cycles of growth (∼64 hrs) using standard conditions^63^. Antibodies were initially screened at 0.5 mg/mL for inhibitory activity before 2-fold serial dilution dose response curves were undertaken to define potency for inhibitory antibodies. Parasitemia was determined using flow cytometry (BD Acurri) after staining with 10 mg/mL of ethidium bromide, with data analysed using FlowJo software (BD Life Sciences). *P. knowlesi* growth in the presence of antibodies was compared to that of untreated control wells to define growth inhibitory activity. All experiments were performed a minimum of three times with duplicate wells unless stated otherwise.

*P. falciparum* growth inhibition assays were performed as described previously^18^.

*P. vivax* growth was analysed using an *ex vivo* invasion assay performed as described previously with slight modification^64^. *P. vivax* samples were collected in 2023 from infected individuals in Kampong Speu, Western Cambodia. *P. vivax* infection was determined using rapid diagnostic testing (CareStartTM Malaria Pf/pan rapid diagnostic tests, Accessbio) or microscopy and species-specific PCR to ensure monoinfection. Venous blood was collected in lithium heparin tubes and immediately sent on ice to the Malaria Research Unit at Institute Pasteur, Cambodia. There, erythrocytes were separated from the plasma, and the plasma was discarded. Erythrocytes were then suspended in warm Roswell Park Memorial Institute (RPMI) medium before leukocyte depletion using a nonwoven fabric filter. The work presented here was approved by the National Ethics Committee for Health Research in Cambodia (192NECHR, July 11, 2022). All patients and/or their parents/guardians provided informed written consent for samples to be taken and used for these purposes.

Infected erythrocytes were enriched using a potassium chloride (KCl)-Percoll density gradient and then transferred into culture in supplemented Iscove′s Modified Dulbecco′s Medium (IMDM)(Gibco) (supplemented with 0.5% (w/v) Albumax II (Gibco), 2.5% (v/v) heat-inactivated human serum, 25 mM HEPES (Gibco), 20 μg/mL gentamicin (Sigma) and 0.2 mM hypoxanthine (C-C Pro)). The stage of the parasite culture was then assessed via thin blood smear. In the case of a majority ring culture, parasites were allowed to mature through to the schizont stage (∼40 hours) before starting the experiment. If the culture consisted mainly of trophozoites, the experiment was carried out after 18-20 hours. The enriched schizonts were then mixed 1:1 with reticulocytes (previously enriched from cord blood or adult peripheral blood from malaria-naïve donors) and pre-labelled with Celltrace Far Red Dye for quantitation. The cultures were incubated with either 500 μg/mL (anti-tetanus toxin 43038) or either 100 μg/mL or 500 μg/mL (monoclonal antibodies and nanobodies) of biologics in a volume of 50 μL in 384-well plates. Cells were stained with Hoechst 33342 to stain parasite DNA and parasitemia was quantified via flow cytometry, with new infections being defined as Far Red/Hoechst double-positive cells. For quantitation, data were normalized against parasites mock treated with PBS. Observed control invasion rates ranged from 0.46 – 5.3% (median = 0.7%).

*P. cynomolgi* assays were performed as previously described^43^. Both *P. falciparum* and *P. knowlesi* GIAs were carried out in parallel to *P. cynomolgi* assays to serve as positive controls for antibody inhibition. *P. cynomolgi* strain Berok R9 was maintained in rhesus red blood cells (Emory Primate Center) at 2% hematocrit in RPMI 1640 with 10% human O+ serum and gassed (1% O₂, 5% CO₂, 94% N₂) at 37°C. The invasion assay was set up with 0.2% hematocrit and 2-3% schizontemia in 30 µl volumes in 384-well plates using antibodies 2D9, 4H10, 5B3, 5B4, IgG control and heparin (positive control) After 12 h of incubation, the parasite DNA was stained with Vybrant™ DyeCycle™ Violet (Invitrogen), and 100,000 cells were analyzed via a Cytek-Northern Lights flow cytometer. Invasion was measured by the percentage of newly parasitized erythrocytes (CellTrace Far Red+/Vybrant™ DyeCycle™ Violet+). Inhibition was assessed relative to control wells without antibodies. Data analysis was done using GraphPad Prism v10.

### Electron microscopy

All electron microscopy was carried out at the Bio21 Ian Holmes Imaging Centre, University of Melbourne.

For negative staining, purified PkPCR^tail^+5B3 Fab (at ∼0.1 mg/mL) was applied to formvar and carbon coated, glow discharged copper grids (300 mesh, ProSciTech). Four microlitres of protein was incubated for one minute, then blotted off, washed twice in water, and then stained with 1% (w/v) uranyl acetate for two minutes before being blotted and dried thoroughly. The grids were then imaged on a Tecnai F30 operating at 200kV. Two-dimensional classification was performed in Cryosparc (v4.4.1)^65^.

For cryo-electron microscopy, freshly purified PkPCR^tail^ and PkPCR^tail^+5B3 Fab at 0.5-1 mg/mL was applied to glow discharged UltrAuFoil (Quantifoil Micro Tools GmbH) grids (300 mesh, R1.2/1.3) or HexAuFoil (Quantifoil Micro Tools GmbH) and then blotted for 5 seconds with a blot force of 7 (UltrAuFoil) or 10 (HexAuFoil) before being plunged into liquid ethane using a Vitrobot Mark IV (Thermo Fisher Scientific) operating at 4°C and 100% humidity. Grids were screened for good ice quality on an FEI Talos Arctica (Thermo Fisher Scientific) operating at 200 kV. Grids showing sufficient thin and amorphous ice were then transferred to an FEI Titan Krios G4(Thermo Fisher Scientific) for data collection. Data were collected using an acceleration voltage of 300 kV and a Falcon 4i detector (Thermo Fisher Scientific) using EPU automation software. Data were collected over three sessions, two for PkPCR^tail^ (from two independent grids) and one for PkPCR^tail^+5B3. Pixel sizes used for collection were 0.506 Å/pixel with a total dose of 50 e^-^/Å^2^ for PkPCR^tail^ and 0.808 Å/pixel with a total dose of 40 e^-^/Å^2^ for PkPCR^tail^+5B3 and all datasets were collected with a nominal defocus range of −0.5 µm to −2 µm. CryoSPARC (v4.4.1-v4.6.2)^65^ was used for all data processing. Gain and motion corrected, and contrast transfer function (CTF)- estimated movie stacks were curated to select for good CTF fit and to remove micrographs that showed signs of significant drift, contained obvious frost contamination, or had no visible particles. This resulted in 8,730 and 826 movie stacks for session one and two, respectively, for PkPCR^tail^ and 3,285 movie stacks for PkPCR^tail^+5B3. The curated datasets were then used in multiple rounds of automated picking and 2D class averaging. For PkPCR^tail^ 11,955 particles corresponded to the ‘front view’, and 3,067 particles corresponded to the ‘side view’. For PkPCR^tail^+5B3 13,478 particles corresponded to the ‘front view’, and 2,694 particles corresponded to the ‘side view’. Whilst clear features were visible in these small number of classes, severe orientation bias and the small, flat, and elongated shape of the particles precluded three-dimensional reconstruction.

### Reticulocyte enrichment for flow cytometric binding assays

Cord blood was obtained through a material transfer agreement (MTA, ID# M19/110) with the Bone Marrow Donor Institute (BMDI) at the Royal Children’s Hospital in Melbourne, Australia under the human ethics project “14/09, Malaria parasite growth and invasion into reticulocytes” which was approved by the Walter and Eliza Hall Institute Human Research Ethics Committee (HREC). Cord blood was passed through a RC High Efficiency Leucocyte Removal Filter (Haemonetics Australia) and then centrifuged at 2000 x *g* for five minutes to separate blood from serum. The blood was then washed three times in 1x human tonicity PBS (HTPBS) before being made up to 50% hematocrit. This 50% solution was then layered on top of a 70% (v/v) Percoll cushion (GE Healthcare). Centrifugation for 25 minutes at 2100 x*g* separated the mature erythrocytes from the reticulocytes, with the reticulocytes forming a thin band at the interface between buffer and Percoll. Reticulocytes were stored in 1x HTPBS at 4°C.

### Flow cytometry-based erythrocyte binding assays

For assays using mature erythrocytes, erythrocytes were washed twice in PBS and then made up to a density of approximately 1 x 10^7^ cells/mL in PBS + 1% (w/v) BSA (PBS-BSA). Each sample used 100 μL of this suspension. Erythrocytes were centrifuged, the supernatant was removed, and the cells were resuspended in a solution containing freshly prepared recombinant proteins in PBS-BSA. Individual proteins were prepared at a final concentration of 2 μM (except for PfRh5, which was prepared at 400 nM), and complexes were mixed with an equimolar amount of protein to a final concentration of 2 μM. After a 45-minute incubation at room temperature, the samples were centrifuged, washed, and then incubated with primary antibodies, either 5A9 (anti-Rh5), 4E2 (anti-PC) or 5E11 (anti-Ripr). After a 45-minute incubation the cells were again centrifuged and then incubated with Alexa-488 anti-mouse fluorescent antibody at a dilution of 1:100. After a 45- minute incubation, cells were washed twice in PBS and then resuspended before analysis on an Attune NxT flow cytometer (Thermo Fisher Scientific). For each sample 50,000 events were recorded. The data were then analysed in FlowJo^TM^ v10.7 Software (BD Life Sciences). Antibody background was subtracted from the positive population recorded in the presence of recombinant protein and this background-subtracted value has been plotted in the summary figures.

For assays involving reticulocyte-enriched cord blood, erythrocytes were made up in 1x HTPBS + 1% (w/v) BSA (HTPBS-BSA) to a density of approximately 1 x 10^7^ cells/mL. Each sample used 100 μL of this suspension. Reticulocytes were centrifuged (2000 x *g* for one minute), the HTPBS- BSA removed, and then resuspended in a solution containing recombinant proteins in HTPBS- BSA and incubated at room temperature for 45 minutes. PvPC and PkPC were used at a final concentration of 2 μM. Samples were centrifuged after which the protein solution removed, and cells were washed once with HTPBS-BSA and then incubated with 4E2 (anti-PC) at a concentration of 0.05 mg/mL or polyclonal sera (anti-RBP2b) at a concentration of 12.5 μg/mL. After a 45-minute incubation, the cells were again centrifuged, and the antibody solution was removed. Cells were washed once as before and then stained with Alexa-647 (either anti-rabbit or anti-mouse) at a dilution of 1:100. After 45 minutes the reticulocytes were again washed and incubated with 50 μL of thiazole orange (BD Retic-Count, BD Biosciences) for 30 minutes. Finally, the reticulocytes were centrifuged, the Retic-Count solution removed, and cells were washed with 1x HTPBS two times before analysis on an Attune NxT flow cytometer (Thermo Fisher Scientific). For each sample 50,000 events were recorded. The data were then analysed in FlowJo^TM^ v10.7 Software (BD Life Sciences). This involved gating reticulocytes and then applying a quadrant gate according to the thiazole orange staining and the background staining of the antibody in combination with the Alexa 647. This antibody background was subtracted from the positive population recorded in the presence of recombinant protein: this background-subtracted value has been plotted in the summary figures. Positive binding is determined by the double positive population in the upper right-hand quadrant.

### Data visualization

All data visualization was done in University of California, San Francisco (UCSF) ChimeraX versions 1.2-1.8 (https://www.cgl.ucsf.edu/chimerax/)^66^. PyMOL was utilised for structure alignment and calculation of root mean square deviation (RMSD) (https://www.pymol.org/)^67^.

### Data availability

The crystal structure reported in this manuscript has been deposited in the Protein Data Bank, www.rcsb.org (PDB ID code 9NSD).

## Extended Data

**Extended Data figure 1.**
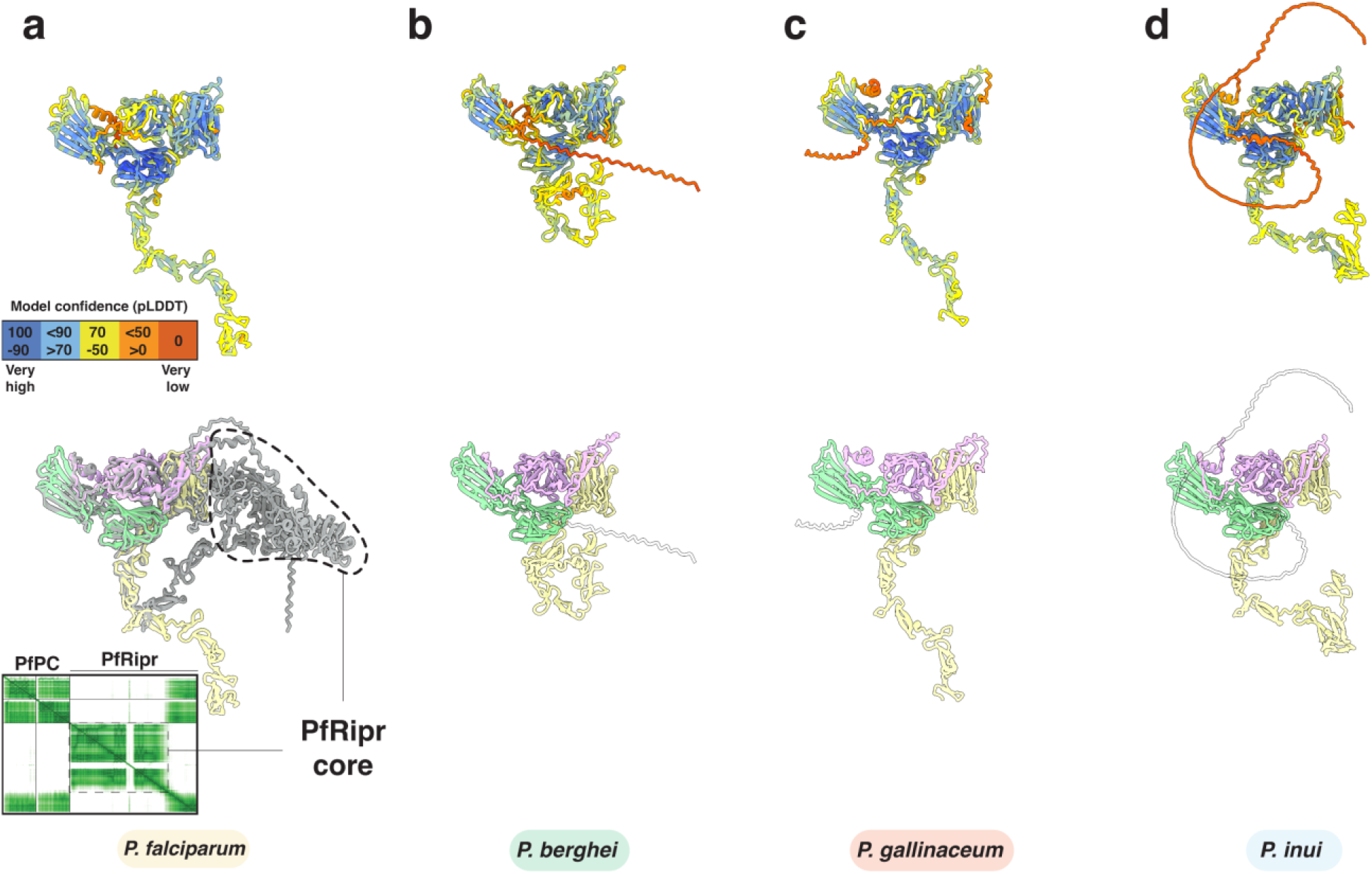
AlphaFold 3 predicts a PTRAMP, CSS, Ripr complex for all major clades of *Plasmodium*. **a-d.** AlphaFold 3 predictions of PTRAMP, CSS and Ripr from several species of *Plasmodium*. Top row: Models colored by confidence (pLDDT). Bottom row: models colored as follows: PTRAMP (pink), CSS (green) and Ripr (yellow). The prediction for *P. falciparum* shows the alignment with full length Ripr, where the core forms an ordered structure. Predicted alignment error (PAE) plot is shown as inset. All models have transmembrane domains and signal peptides omitted for clarity.

**Extended Data figure 2.**
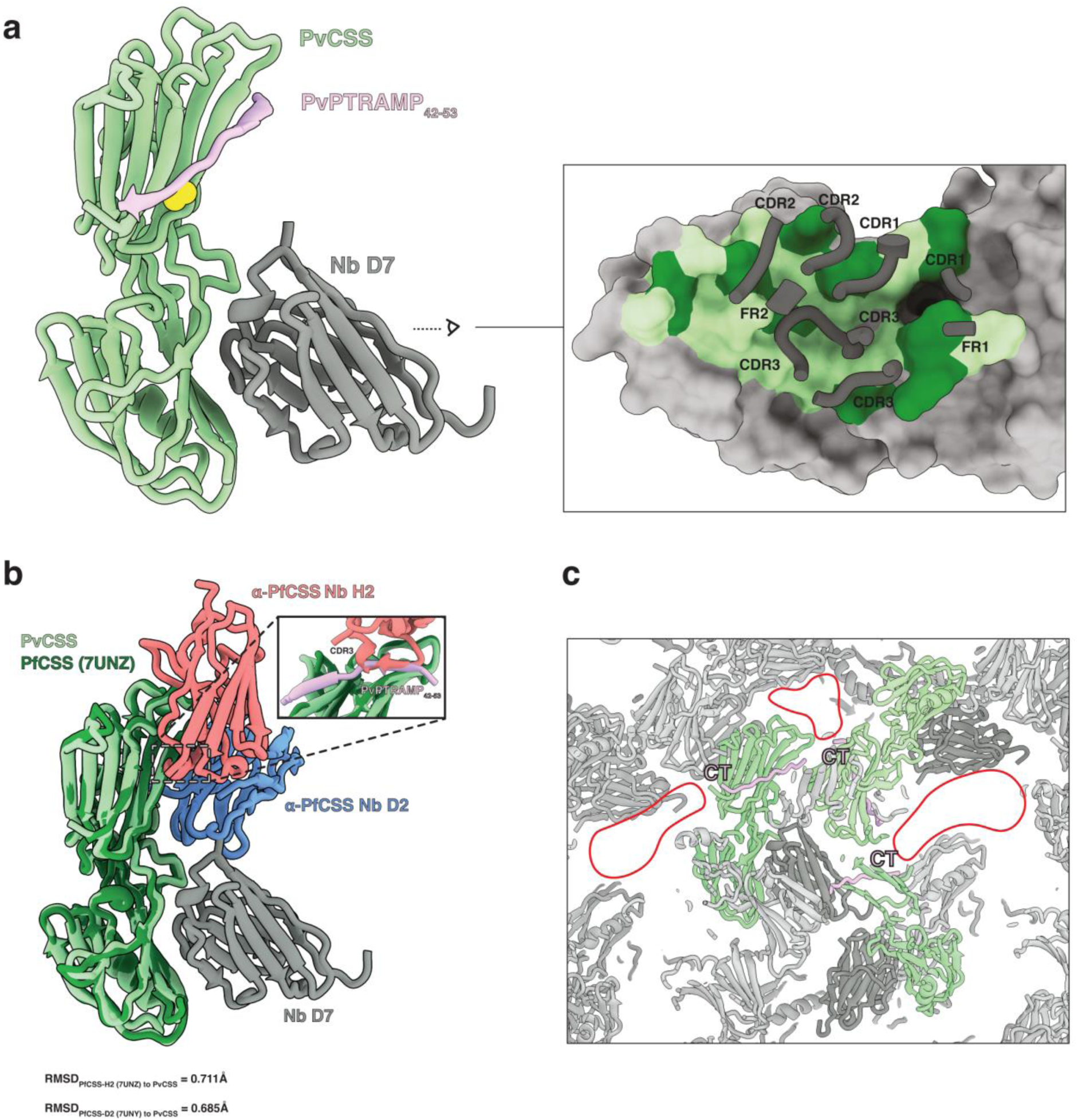
Comparison of PvCSS crystal structure with PfCSS structures. **a.** Binding interface between nanobody D7 and PvCSS. The detail shows the entire interface. PvCSS is grey and the region of D7 binding is shown in pale green, with hydrogen bonds and salt bridges shown in forest green. Regions of D7 involved in the interaction are shown and consist of complementarity determining regions (CDR) and framework regions (FR). Regions of D7 not involved in binding are omitted for clarity. **b.** Overlay of PfCSS-H2 (H2 in red) and PfCSS-D2 (D2 in blue) crystal structures with PvCSS-PvPTRAMP-D7(D7 in grey) crystal structure. Inset shows PvPTRAMP_42-53_ occupies the same region as the CDR3 loop of H2, explaining how it competes with PfPTRAMP binding to PfCSS^1^. **c.** Crystal packing of PvCSS-PvPTRAMP-D7. The asymmetric unit is shown in color (PvCSS in green, PvPTRAMP in pink and D7 in dark grey) and other symmetry related copies in light grey. Red circles show pockets within the crystal lattice that are adjacent to the C-terminal (CT) of PvPTRAMP_42-53_ that likely accommodate the GFD and TSR domains of PvPTRAMP.

**Extended Data figure 3.**
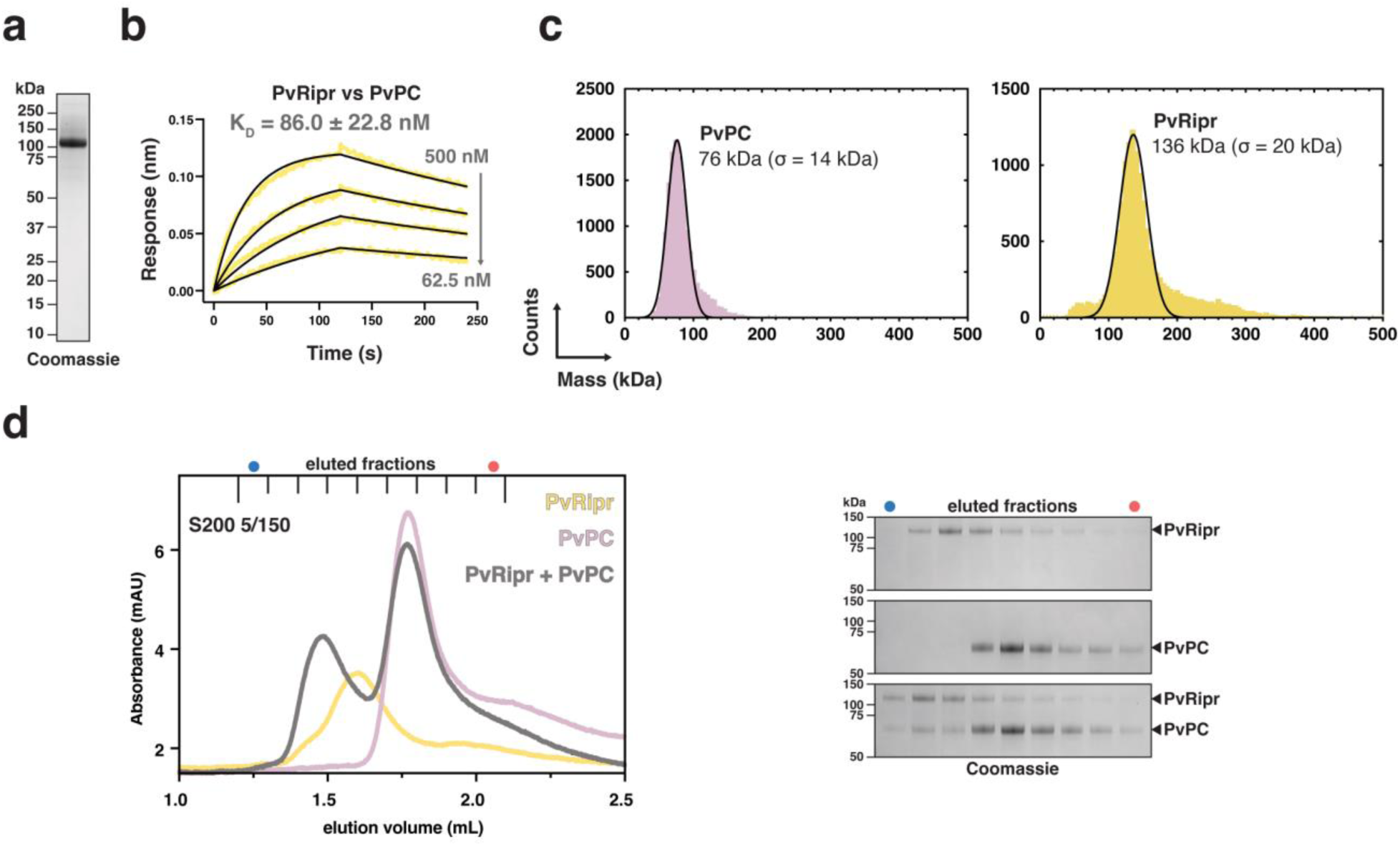
The PvPCR complex can be reconstituted *in vitro* using recombinant proteins. **a.** Representative SDS-PAGE of recombinant full-length PvRipr. **b.** Representative biolayer interferometry sensorgram of PvRipr vs PvPC. Data are in yellow and 1:1 model best fit in black. **c.** Mass distribution plots of PvPC (pink) and PvRipr (yellow) as determined by mass photometry. Histogram data are in color and the Gaussian curve fit in black. **d.** Size-exclusion chromatography of PvPC (pink), PvRipr (yellow) and PvPCR (grey), with SDS-PAGE results of the co-complexation, showing co-elution of PvPC and PvRipr.

**Extended Data figure 4.**
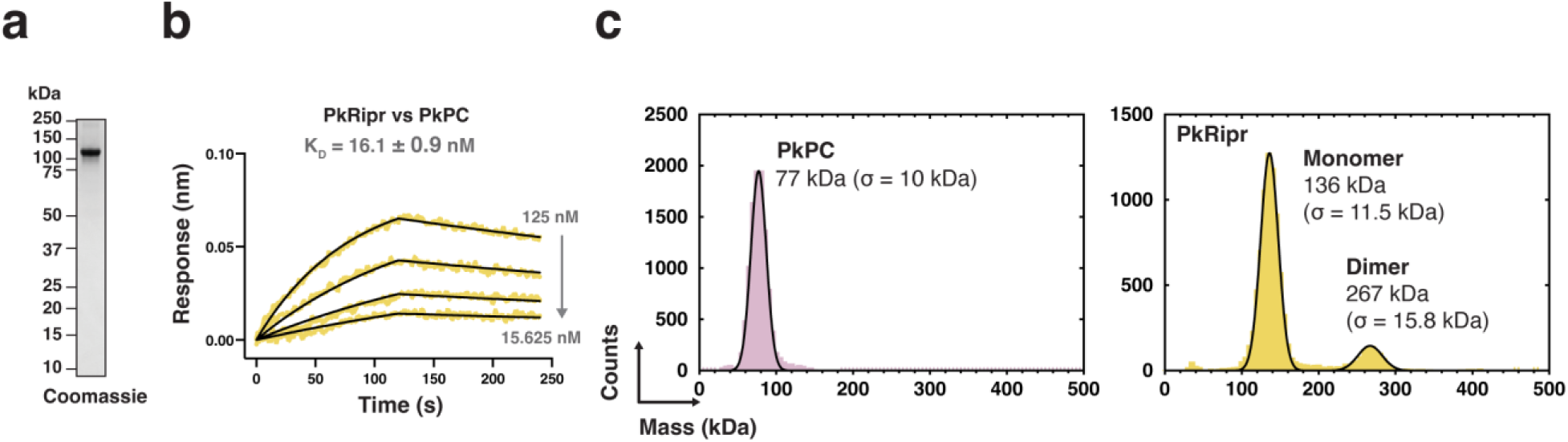
Biophysical analysis of PkPC and PkRipr. **a.** SDS-PAGE of recombinant full-length PkRipr. **b.** Representative biolayer interferometry sensorgram of PkRipr vs PkPC. Data are in yellow and 1:1 model best fit in black. **c.** Mass distribution plots of PkPC (pink) and PkRipr (yellow) as determined by mass photometry. Histogram data are in color and the Gaussian curve fit in black.

**Extended Data figure 5.**
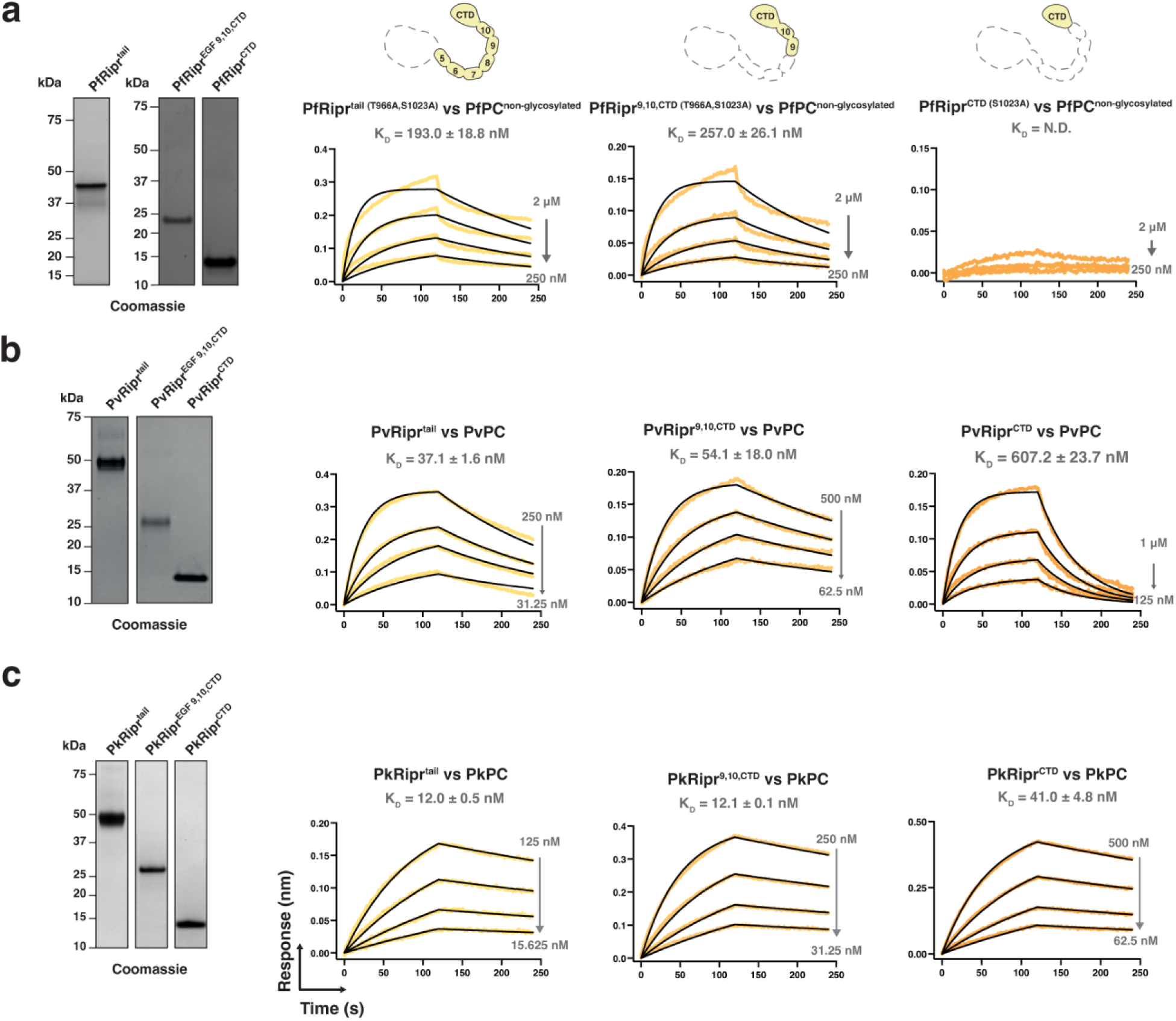
Biophysical analysis of Ripr truncations. **a.** SDS-PAGE of recombinant PfRipr truncations and representative biolayer interferometry sensorgrams of Ripr truncations vs PfPC. PfRipr^tail^ and PfRipr^EGF 9,10,CTD^ harbor T966A and S1023A mutations and PfRipr^CTD^ harbors S1023A mutation. **b.** SDS-PAGE of recombinant PvRipr truncations and representative biolayer interferometry sensorgrams of Ripr truncations vs PvPC. **c.** SDS-PAGE of recombinant PkRipr truncations and representative biolayer interferometry sensorgrams of Ripr truncations vs PkPC. For all sensorgrams, data are in color and 1:1 model best fit in black where applicable. Schematics of the Ripr truncations are shown above.

**Extended Data figure 6.**
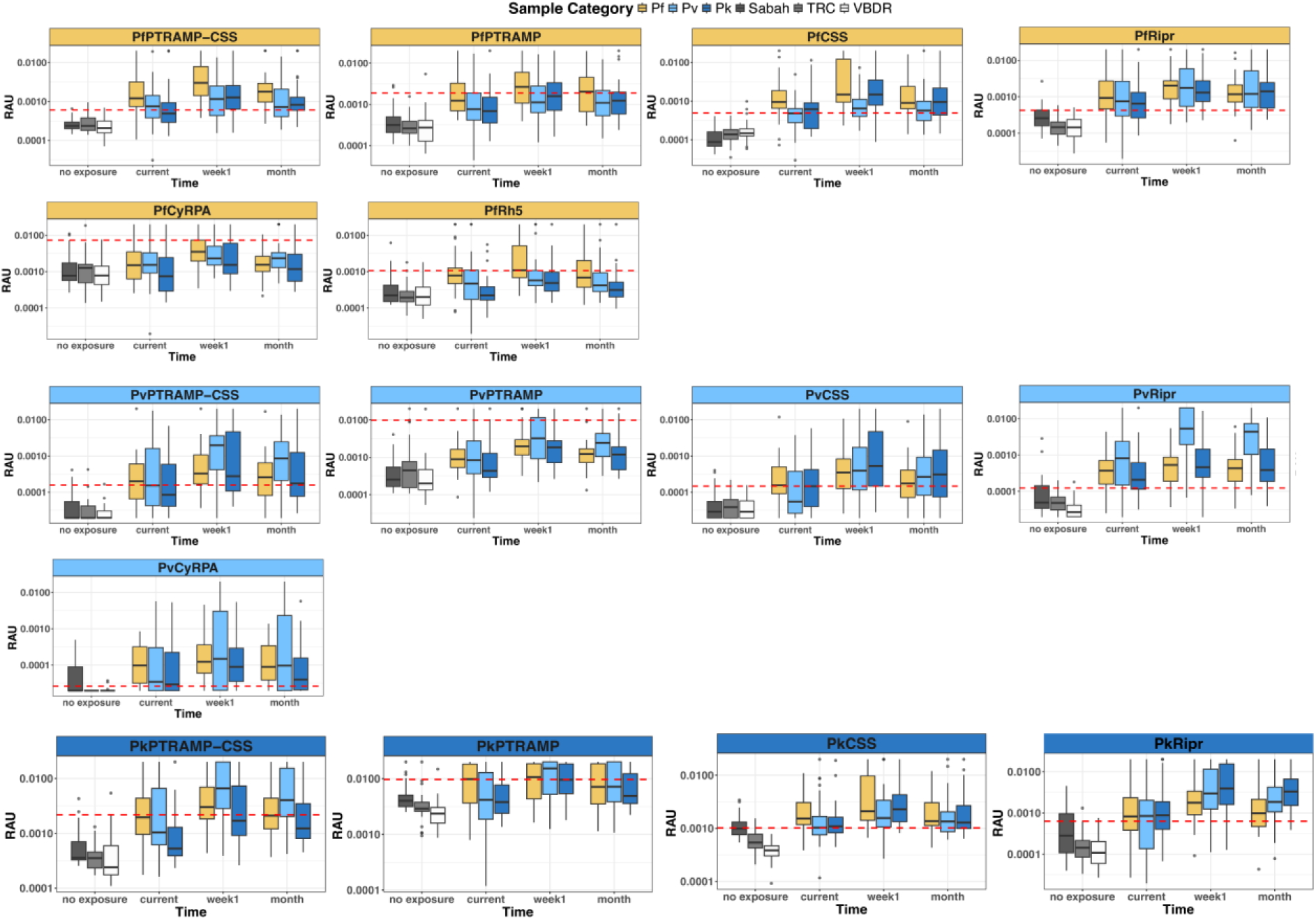
Serological assessment of invasion antigens. Antibody kinetics in longitudinal plasma samples collected from individuals with *P. falciparum* (yellow), *P. vivax* (light blue), *P. knowlesi* (dark blue) infections at three timepoints (current (n=95), week 1 (n=98) and a month (n=98)) on a logarithmic scale. Samples from Volunteer Biospecimen Donor Registry (white, n =28) and Thailand Red Cross (light grey, n = 29) was used to set the sero-positivity cut-off (red dotted line), which is mean of these samples + 2 x standard deviation. Samples from healthy (non-febrile) individuals from Sabah Malaysia (dark grey, n=30) are not included in the sero-positivity calculation as their previous exposure status to any of the *Plasmodium* species is unknown. Boxplots display the median (horizontal line), interquartile range (IQR) (the box), largest and smallest values (1.25 x IQR, whiskers) and outliers are displayed as points.

**Extended Data figure 7.**
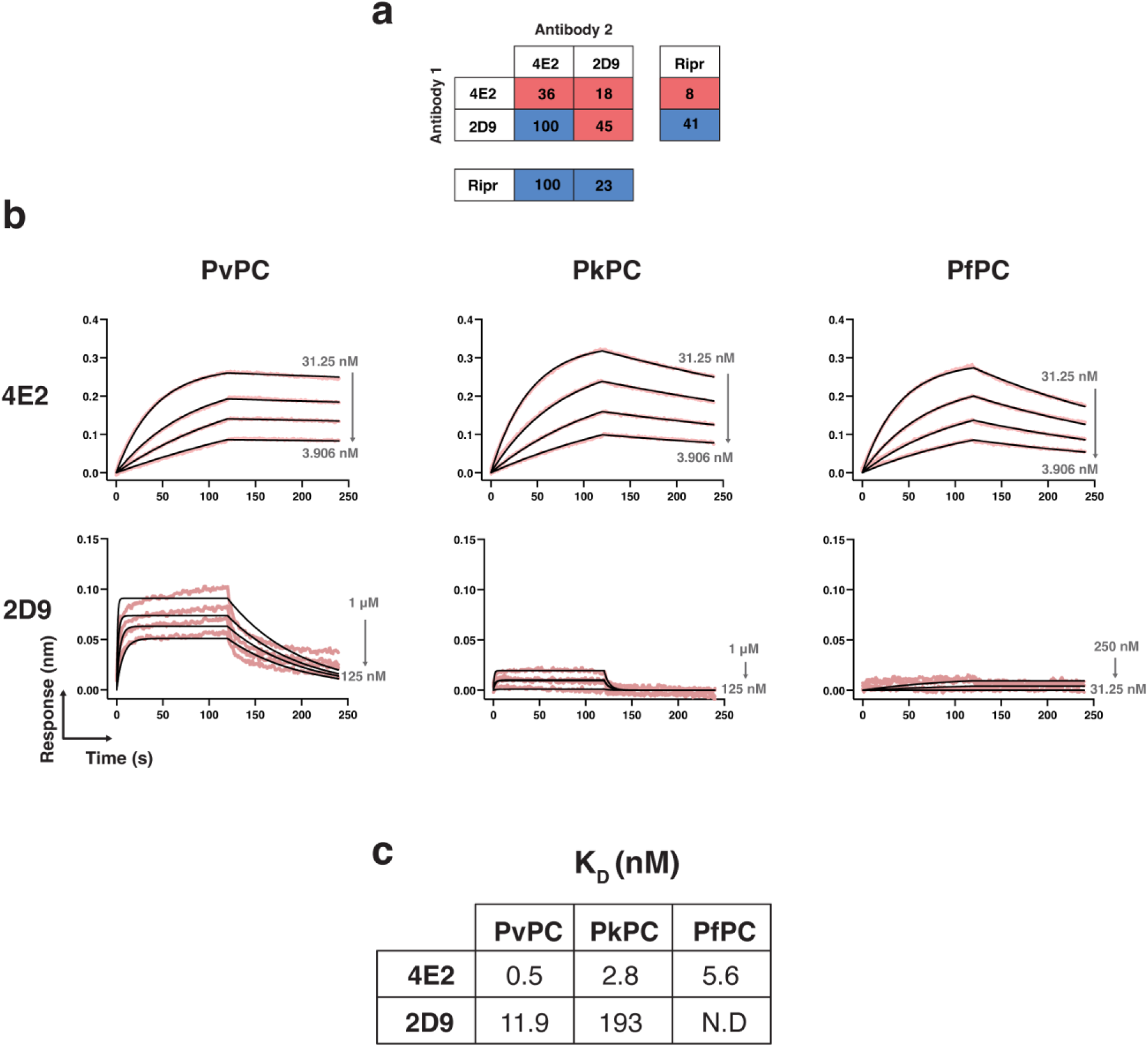
Characterization of anti-PvPC mouse monoclonal antibodies. **a.** Competition binning of anti-PvPC monoclonal antibodies and PvRipr. **b.** Representative biolayer interferometry sensorgrams of antibody binding to PvPC, PkPC, and PfPC. Data are in color and 1:1 model best fit in black. **c.** Table of K_D_ values, in nM, for the curves in b. N.D = not determined

**Extended Data figure 8.**
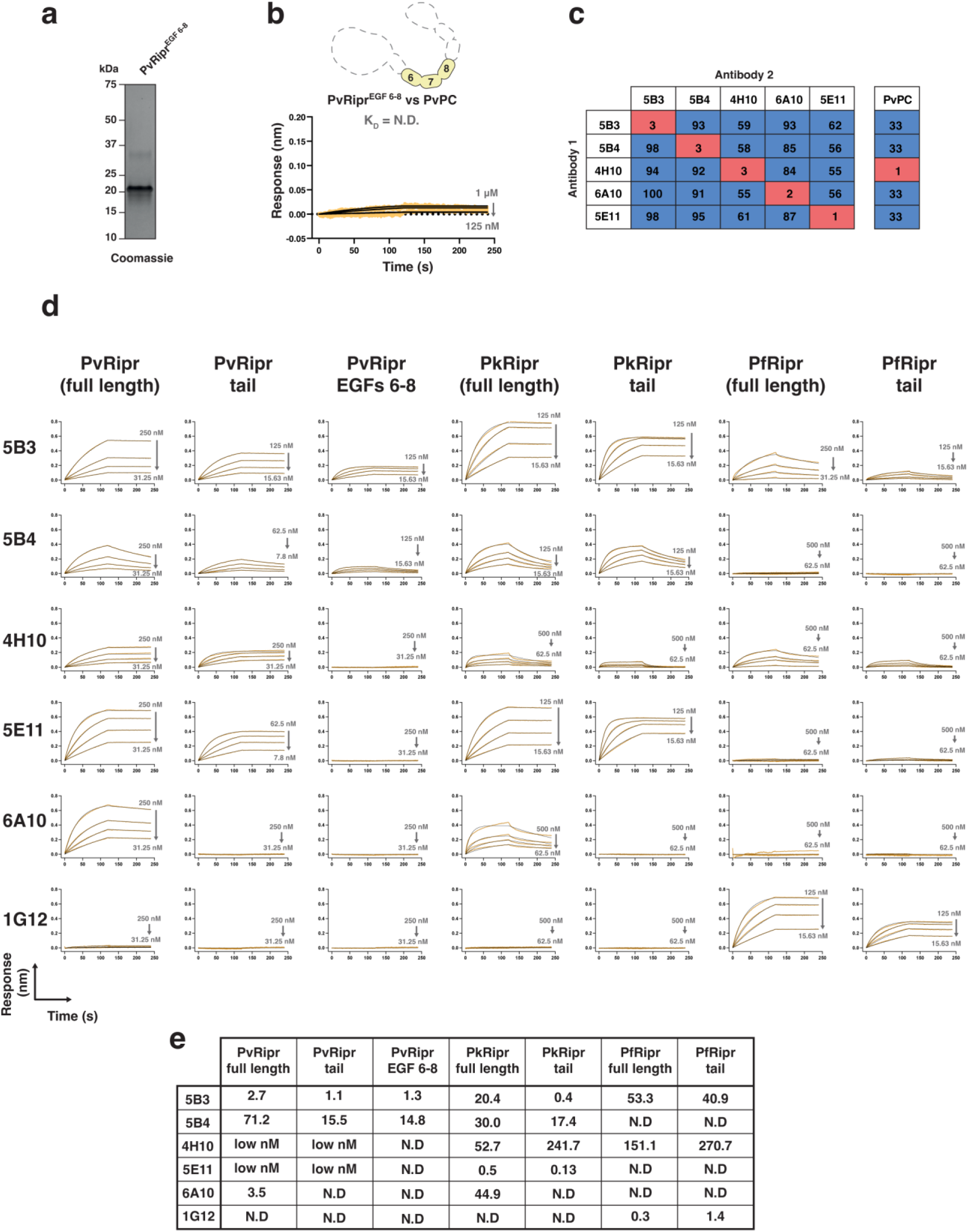
Characterization of anti-PvRipr mouse monoclonal antibodies. **a.** SDS-PAGE of recombinant PvRipr^EGF6–8^. **b.** Biolayer interferometry sensorgram of PvRipr^EGF6–8^ vs PvPC showing no interaction between the two proteins. **c.** Competition binning of anti-PvRipr antibodies. **d.** Representative biolayer interferometry sensorgrams of antibody binding to PvRipr, PkRipr and PfRipr. The anti-PfRipr antibody 1G12 was included as a control ^2^. Data are in color and 1:1 model best fit in black. **e.** Table of K_D_ values, in nM, for the curves in d. N.D. = not determined.

**Extended Data figure 9.**
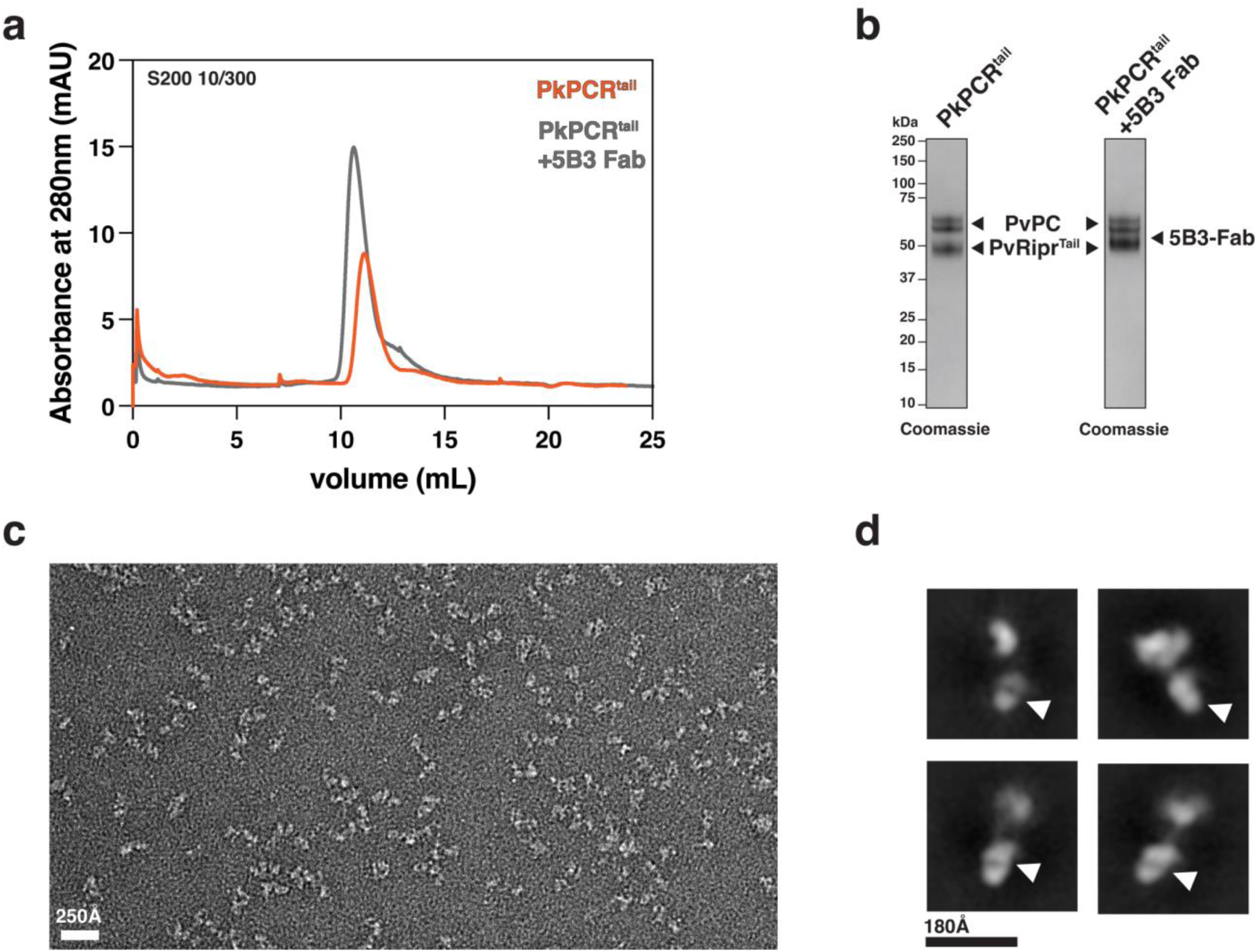
Negative stain electron microscopy of PkPCR^tail^+5B3 Fab shows an elongated structure. **a.** Size-exclusion chromatography chromatograph of PkPCR^tail^ and PkPCR^tail^+5B3 Fab. **b.** Non-reducing SDS-PAGE of peak fractions of purified complexes from a. 5B3 Fab and Ripr^tail^ have approximately the same molecular weight. **c.** Representative electron micrograph of negatively stained PkPCR^tail^+5B3 Fab particles. **d.** Two-dimensional class averages of PkPCR^tail^+5B3 Fab. White arrows show density attributable to 5B3-Fab.

**Extended Data figure 10.**
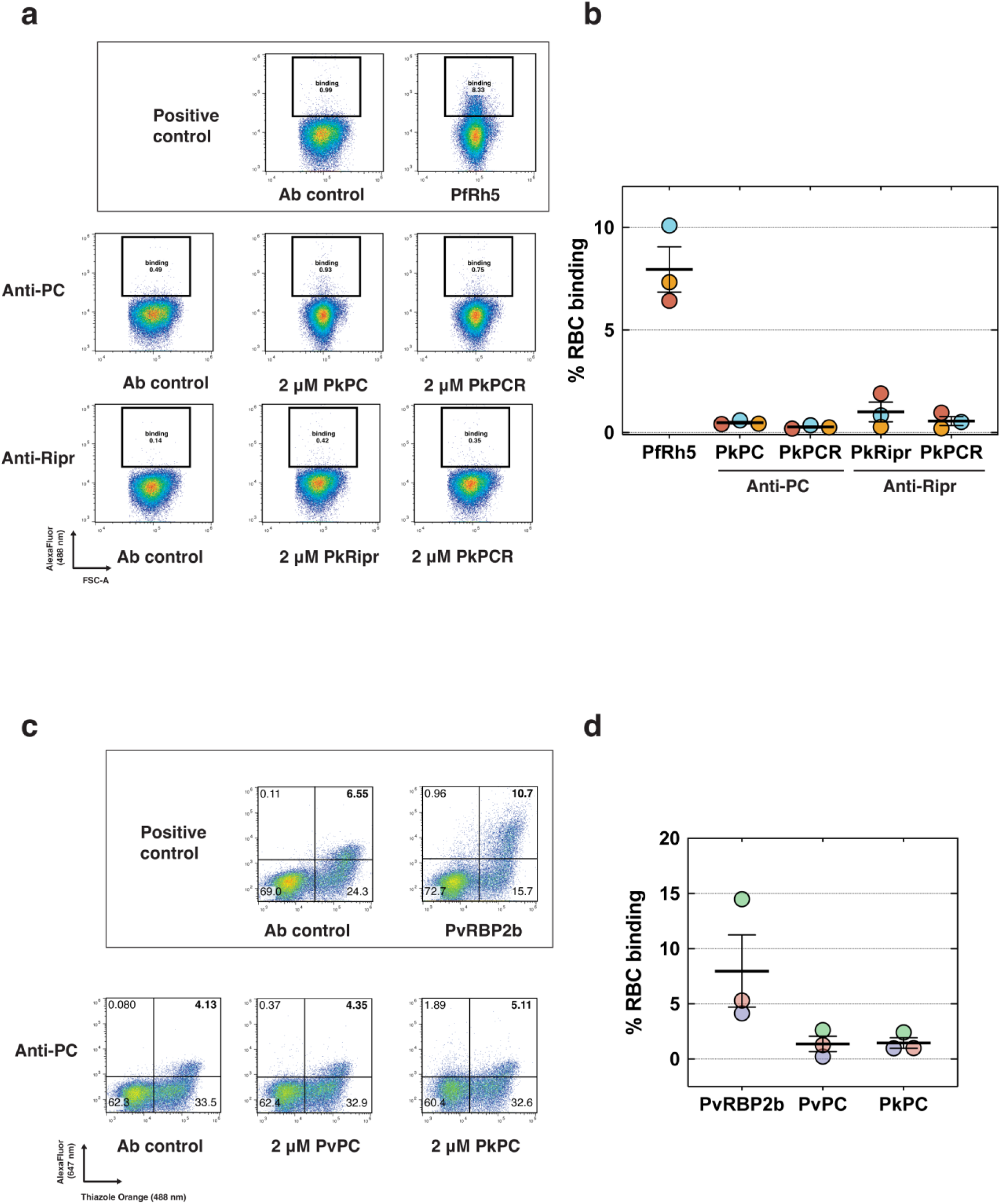
PvPC, PkPC and PkPCR do not bind erythrocytes. **a.** Representative flow cytometry plots of PkPC, PkRipr, and PkPCR incubated with erythrocytes and detected with either anti-PC or anti-Ripr antibodies. PfRh5 was used as a positive control for erythrocyte binding. **b.** Quantitation of binding events from three independent experiments. Individual experimental replicates are depicted as colored dots, and the mean is shown with standard error of the mean (SEM). **c.** Representative flow cytometry plots of PvPC and PkPC incubated with reticulocyte enriched cord blood and detected with anti-PC antibody. PvRBP2b was used as a positive control for reticulocyte binding. **d.** Quantitation double positive binding events of three independent experiments. Individual colored dots represent a replicate performed with a single donor’s blood. Mean is shown with SEM.

