## Supplementary Information for "PTRAMP, CSS and Ripr form a conserved complex required for merozoite invasion of *Plasmodium* species into erythrocytes"

**Table S1. Data collection and refinement statistics of PvCSS-PvPTRAMP-D7 structure**

| | **PvCSS_PvPTRAMP_D7** |
|---|---|
| **Beamline** | MX2 |
| **Wavelength** | 0.953739 |
| **Resolution range**[^] | 48.32 - 3.14 (3.252 - 3.14) |
| **Space group** | P 32 |
| **Cell dimensions** | |
| **a,b,c (Å)** | 125.2, 125.2, 106.5 |
| **α, β, γ (°)** | 90, 90, 120 |
| **Total reflections** | 174, 965 (17,387) |
| **Unique reflections** | 32, 570 (3,251) |
| **Multiplicity** | 5.4 (5.4) |
| **Completeness (%)** | 99.9 (99.9) |
| **<I/σ I>** | 13.2 (1.5) |
| **$R_{merge}$ (%)** | 9.6 (104.5) |
| **$R_{pim}$ (%)** | 4.6 (49.7) |
| **$CC_{½}$ (%)** | 99.8 (54.4) |
| **Reflections used in refinement** | 32, 565 (3, 250) |
| **Reflections used for R-free** | 1, 700 (215) |
| **$R_{work}$ (%)** | 22.5 (35.4) |
| **$R_{free}$ (%)**[&] | 27.3 (37.3) |
| **Number of non-hydrogen atoms** | 9, 551 |
| **macromolecules** | 9, 551 |
| **Protein residues** | 1, 196 |
| **RMS (bonds) (Å)** | 0.003 |
| **RMS (angles) (°)** | 0.67 |
| **Ramachandran plot** | |
| **Ramachandran favored (%)** | 92.2 |
| **Ramachandran allowed (%)** | 7.8 |
| **Ramachandran outliers (%)** | 0.0 |
| **Rotamer outliers (%)** | 1.03 |
| **Clashscore** | 7.3 |
| **Wilson B-factor** | 97.5 |
| **Average B-factor** | 93.6 |

[^] Statistics for the highest-resolution shell are shown in parentheses.
[&] 5% of data were used for $R_{free}$ calculation

**Table S2. Table of contacts between PvCSS and PvPTRAMP**

| **PvCSS Residue (BSA Å$^{2}$)** | **Interaction type** | **PvPTRAMP residue** |
|---|---|---|
| **Arg118 (23.8)** | | |
| Arg | VDW | Gly53 |
| Arg$^{N}$ | HB | Gly53$^{O}$ |
| **Ala119 (21.2)** | | |
| Ala | VDW | Val52, Gly53 |
| Ala$^{N}$ | HB | Gly53$^{O}$ |
| Ala$^{O}$ | HB | Gy53$^{N}$ |
| **Asn120 (26.0)** | | |
| Asn | VDW | Ala51, Val52, Gly53 |
| **Leu121 (52.9)** | | |
| Leu | VDW | Cys50, Ala51, Val52, Gly53 |
| Leu$^{N}$ | HB | Ala51$^{O}$ |
| Leu$^{O}$ | HB | Ala51$^{N}$ |
| **Cys122 (38.0)** | | |
| Cys | VDW | Glu49, Cys50, Ala51 |
| Cys$^{S\gamma}$ | DSB | Cys50$^{S\gamma}$ |
| **Ser123 (41.1)** | | |
| Ser | VDW | Pro48, Glu49, Cys50 |
| Ser$^{N}$ | HB | Glu49$^{O}$ |
| **Cys124 (3.7)** | | |
| Cys | VDW | Glu47, Glu49 |
| **Phe126 (6.3)** | | |
| Phe | VDW | Glu47 |
| **Arg129 (42.2)** | | |
| Arg | VDW | Leu46, Glu47 |
| **Glu136 (26.9)** | | |
| Glu | VDW | Ile43 |
| **Lys137 (62.1)** | | |
| Lys | VDW | Lys42, Ile43 |
| Lys$^{O}$ | HB | Ile43$^{N}$ |
| **Thr138 (27.9)** | | |
| Thr | VDW | Ile43 |
| **Lys139 (77.0)** | | |
| Lys | VDW | Lys42, Ile43, Val44, Thr45 |
| Lys$^{N}$ | HB | Ile43$^{O}$ |
| Lys$^{O}$ | HB | Thr45$^{N}$ |
| **Val140 (35.8)** | | |
| Val | VDW | Ile43, Thr45, Glu47 |

| | | |
|---|---|---|
| **Val141 (57.8)** | | |
| Val | VDW | Val44, Thr45, Leu46, Glu47 |
| Val$^{N}$ | HB | Thr45$^{O}$ |
| Val$^{O}$ | HB | Glu47$^{N}$ |
| **Cys142 (34.2)** | | |
| Cys | VDW | Glu47, Pro48 |
| **Asn143 (21.5)** | | |
| Asn | VDW | Pro48 |
| **Leu144 (4.0)** | | |
| Leu | VDW | Glu49 |

**Table S3. Table of contacts between PvCSS and nanobody D7**

| PvCSS Residue (BSA Å$^2$) | Interaction type | Nb_D7 residue^ |
|---|---|---|
| **Glu245 (41.9)** | | |
| Glu | VDW | Arg31 |
| Glu$^{O}$ | HB | Arg31$^{N\varepsilon}$ |
| **Val246 (17.1)** | | |
| Val | VDW | Arg31 |
| **Gly247 (33.2)** | | |
| Gly | VDW | Arg31, Tyr32 |
| **Glu248 (94.0)** | | |
| Glu | VDW | Arg31, Ala33, Asn52, Ser52A, Phe98 |
| Glu$^{N}$ | HB | Arg31$^{O}$ |
| Glu$^{O\varepsilon2}$ | HB | Ser52A$^{N}$, Ser52A$^{O\gamma}$ |
| **Tyr250 (64.1)** | | |
| Tyr | VDW | Asn52, Asp53, Phe56 |
| Tyr$^{OH}$ | HB | Asn52$^{N\delta2}$, Asp53$^{O\delta2}$ |
| **Tyr278 (3.5)** | | |
| Tyr | VDW | Tyr102 |
| **Lys279 (66.00)** | | |
| Lys | VDW | Lys94, Gln96, Tyr100, Asp101, Tyr102 |
| Lys$^{N\zeta}$ | SB | Asp101$^{O\delta1}$ |
| **His280 (121.3)** | | |
| His | VDW | Val2, Phe27, Tyr32, Lys94, Asp101, Tyr102 |
| His$^{N}$ | HB | Tyr102$^{OH}$ |
| His$^{O}$ | HB | Lys94$^{N\zeta}$ |
| His$^{N\delta1}$ | HB | Tyr32$^{OH}$ |
| **Asp281 (10.5)** | | |
| Asp | VDW | Tyr32, Lys94 |
| Asp$^{O\delta1}$ | HB | Tyr32$^{OH}$ |
| **Ser283 (37.9)** | | |
| Ser | VDW | Lys94, Gln96, Asp101 |
| **Ser285 (19.7)** | | |
| Ser | VDW | Gln96, Ala97, Phe98 |
| **Ile287 (21.6)** | | |
| Ile | VDW | Phe98 |
| **Leu289 (23.2)** | | |
| Leu | VDW | Phe56 |
| **Lys349 (4.6)** | | |
| Lys | VDW | Tyr58 |

| | | |
|---|---|---|
| **Asn350 (13.5)** | | |
| Asn | VDW | Tyr58 |
| Asn$^{O}$ | HB | Tyr58$^{OH}$ |
| **Phe351 (8.8)** | | |
| Phe | VDW | Tyr58 |
| Phe$^{N}$ | HB | Tyr58$^{OH}$ |
| **Asn352 (83.2)** | | |
| Asn | VDW | Asp50. Tyr58, Phe98, Gly99 |
| Asn$^{N\delta2}$ | HB | Asp50$^{O\delta2}$ |
| **Ala354 (43.6)** | | |
| Ala | VDW | Phe98, Gly99, Tyr100 |
| **Cys355 (1.7)** | | |
| Cys | VDW | Tyr100 |
| **Ala356 (13.8)** | | |
| Ala | VDW | Tyr100 |
| **Lys372 (36.3)** | | |
| Lys | VDW | Gln96, Tyr100 |
| Lys$^{N\zeta}$ | HB | Gln96$^{O\varepsilon1}$, Tyr100$^{OH}$ |
| **Ile374 (41.6)** | | |
| Ile | VDW | Gln96, Ala97, Tyr100 |
| **Thr376 (16.1)** | | |
| Thr | VDW | Ala97, Phe98 |
| Thr$^{O\gamma1}$ | HB | Phe98$^{O}$ |
| **Tyr378 (57.2)** | | |
| Tyr | VDW | Phe56, Thr57, Tyr58, Phe98 |
| Tyr$^{OH}$ | HB | Thr57$^{O}$ |
| **Phe379 (0.6)** | | |
| Phe | VDW | Tyr58 |
| **Asn380 (13.4)** | | |
| Asn | VDW | Thr57, Tyr58 |

^D7 Residues are labelled according to the Kabat numbering system

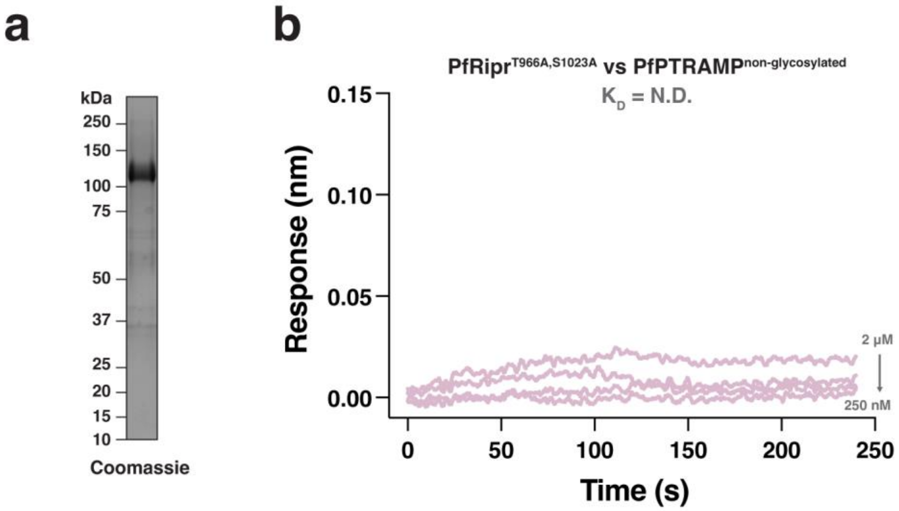

**Supplementary figure 1. Non-glycosylated forms of monomeric PfPTRAMP and PfRipr do not interact with one another. a.** SDS-PAGE of purified recombinant PfRipr$^{T966A,S1023A}$ **b.** Biolayer interferometry sensorgram of PfRipr$^{T966A,S1023A}$ vs PfPTRAMP$^{non\text{-}glycosylated}$ .

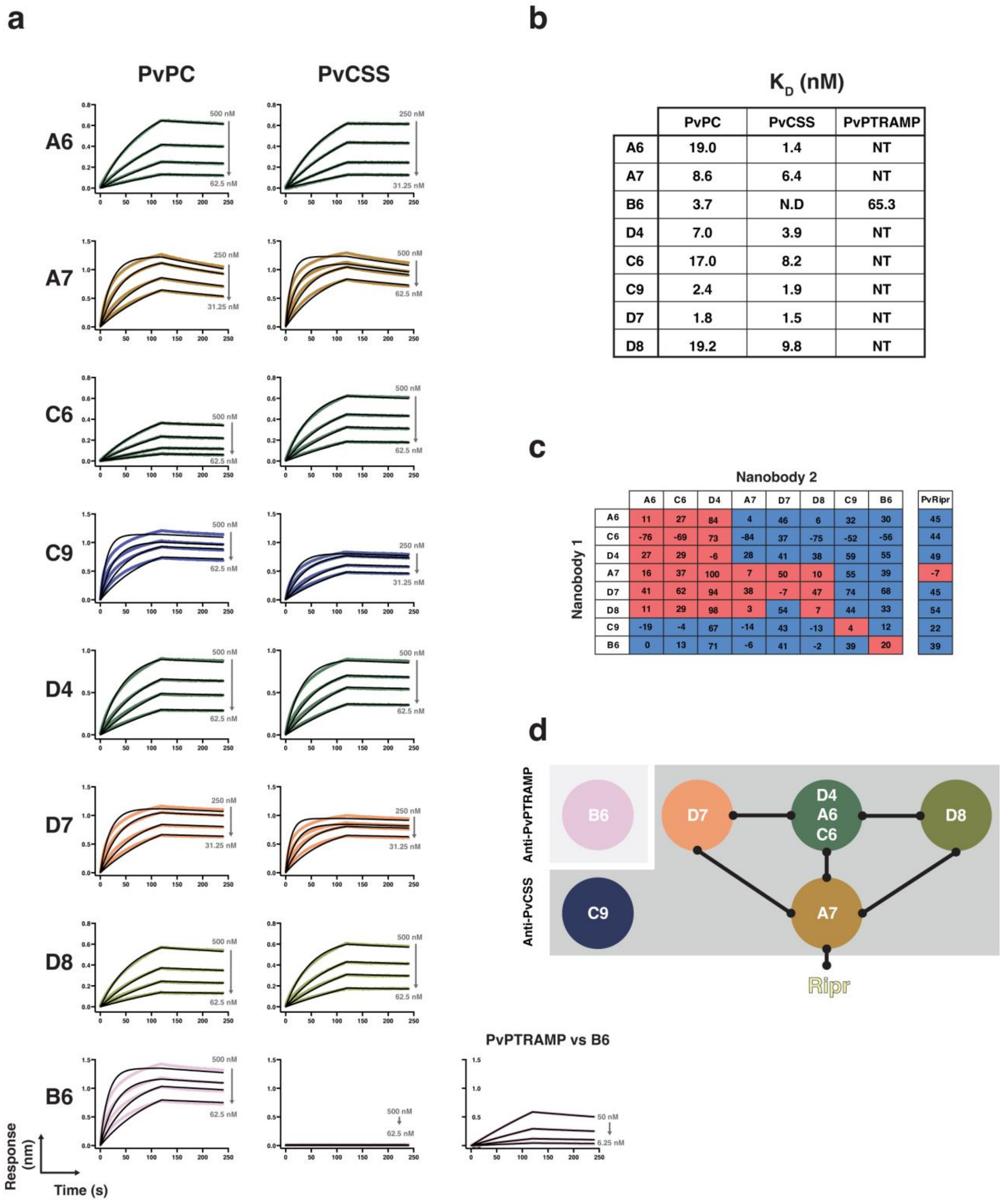

$K_D$ (nM)

| | PvPC | PvCSS | PvPTRAMP |
|---|---|---|---|
| A6 | 19.0 | 1.4 | NT |
| A7 | 8.6 | 6.4 | NT |
| B6 | 3.7 | N.D | 65.3 |
| D4 | 7.0 | 3.9 | NT |
| C6 | 17.0 | 8.2 | NT |
| C9 | 2.4 | 1.9 | NT |
| D7 | 1.8 | 1.5 | NT |
| D8 | 19.2 | 9.8 | NT |

| Nanobody 1 \ Nanobody 2 | A6 | C6 | D4 | A7 | D7 | D8 | C9 | B6 | PvRipr |
|---|---|---|---|---|---|---|---|---|---|
| A6 | 11 | 27 | 84 | 4 | 46 | 6 | 32 | 30 | 45 |
| C6 | -76 | -69 | 73 | -84 | 37 | -75 | -52 | -56 | 44 |
| D4 | 27 | 29 | -6 | 28 | 41 | 38 | 59 | 55 | 49 |
| A7 | 16 | 37 | 100 | 7 | 50 | 10 | 55 | 39 | -7 |
| D7 | 41 | 62 | 94 | 38 | -7 | 47 | 74 | 68 | 45 |
| D8 | 11 | 29 | 98 | 3 | 54 | 7 | 44 | 33 | 54 |
| C9 | -19 | -4 | 67 | -14 | 43 | -13 | 4 | 12 | 22 |
| B6 | 0 | 13 | 71 | -6 | 41 | -2 | 39 | 20 | 39 |

**Supplementary figure 2. Characterisation of anti-PvPC nanobodies. a.** Biolayer interferometry sensorgrams of nanobodies binding to PvPC, PvCSS or PvPTRAMP. Data are shown in color and the 1:1 model best fit shown in black. **b.** Table of $K_D$ values for nanobodies against PvPC and PvCSS. N.D = not determined, NT= not tested. **c-d.** Epitope binning of nanobodies. Boxes are colored on a sliding scale where red represents a competing nanobody and blue represents no competition. Epitope bins from competition with PvRipr are represented in (D). The competition table in (C) has been normalized to the greatest response, which was arbitrarily set to 100. Nanobodies that compete with one another or with Ripr binding are connected with black bars.

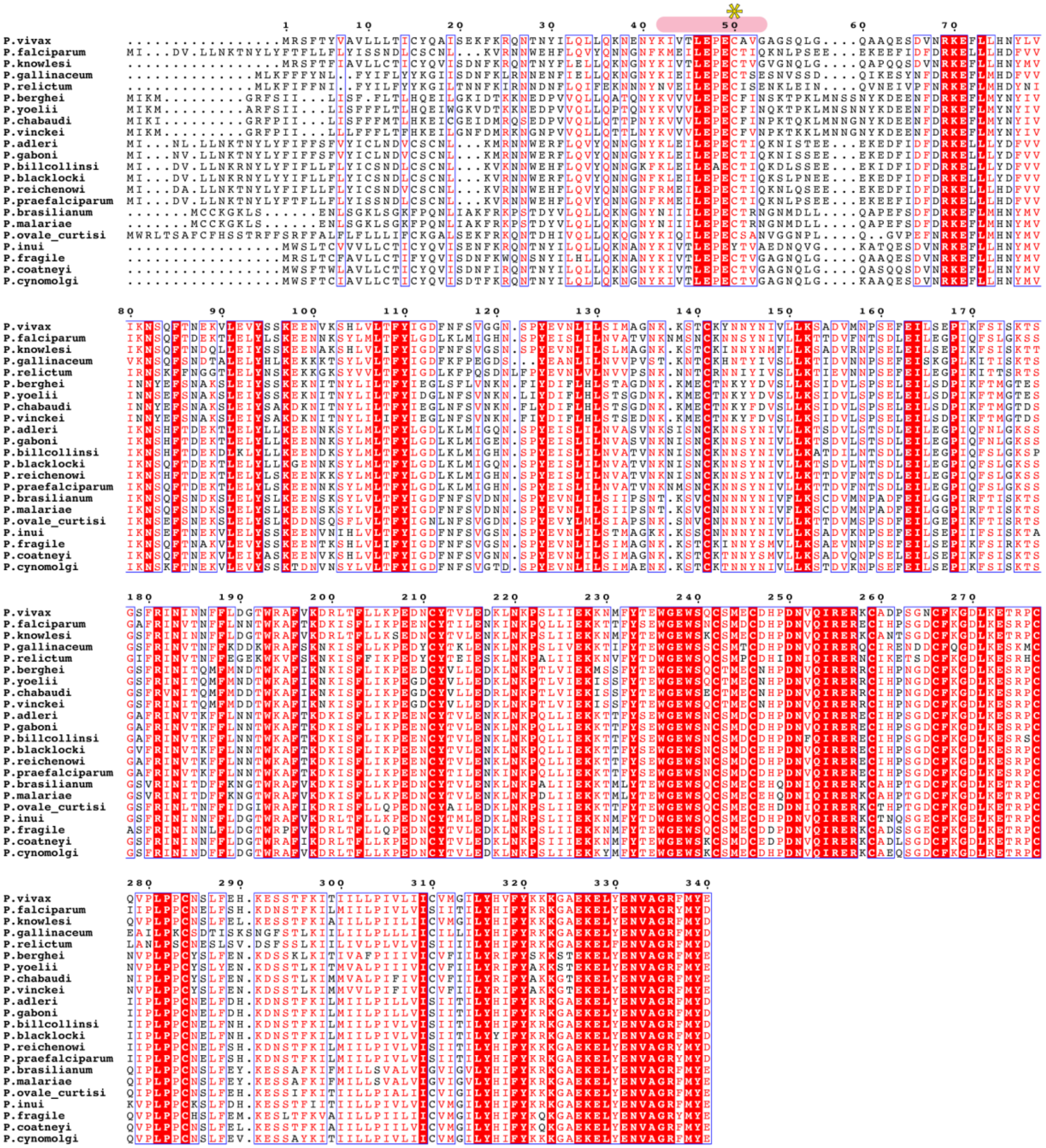

**Supplementary figure 3. Comparison of PTRAMP sequences from a selection of *Plasmodium* species.** Multiple sequence alignment of PTRAMP from several *Plasmodium* species. Regions involved in PvCSS binding are indicated with pink above the interacting amino acids. The unpaired cysteine involved in intermolecular disulfide formation is marked with a yellow asterisk.

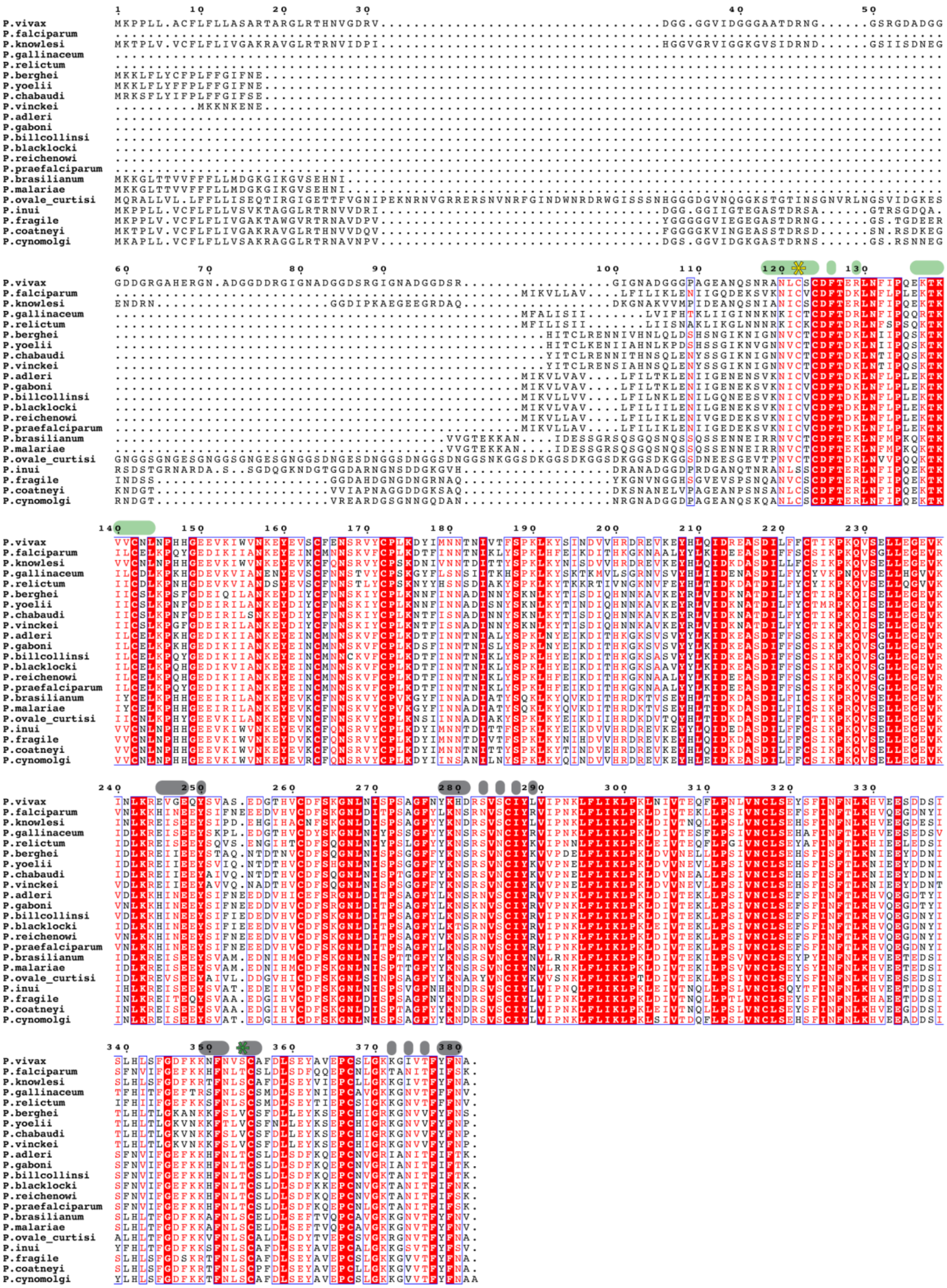

**Supplementary figure 4. Comparison of CSS sequences from a selection of Plasmodium species.** Multiple sequence alignment of CSS from several Plasmodium species. Regions involved in PvPTRAMP binding are indicated by green above the interacting amino acids. Regions involved in the D7 interface are marked grey. The unpaired cysteine involved in intermolecular disulfide formation is marked with a yellow asterisk. The Ser354 that was mutated to remove a potential N-linked glycan is marked by a green asterisk.

a

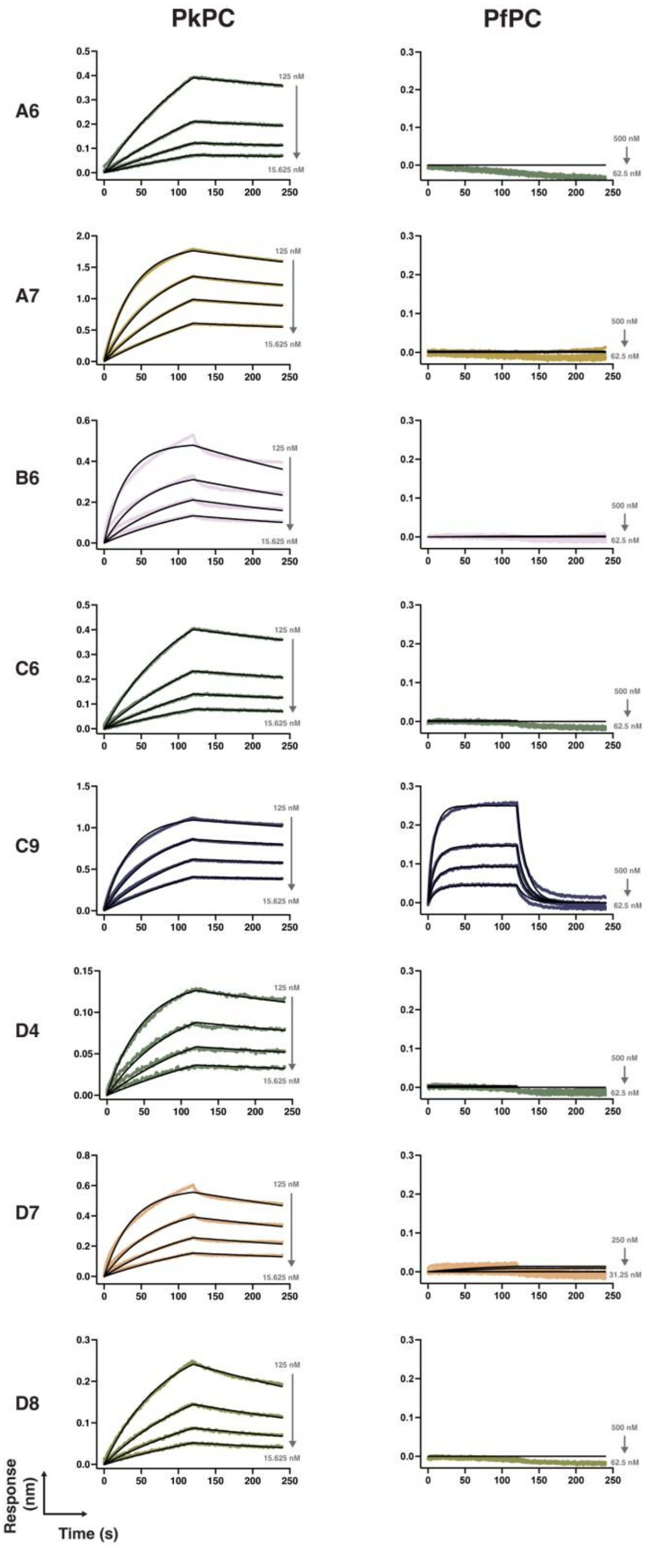

b

$K_D$ (nM)

| | PkPC | PfPC |
|---|---|---|
| **A6** | 14 | N.D |
| **A7** | 4.4 | N.D |
| **B6** | 9.0 | N.D |
| **C6** | 13.8 | N.D |
| **C9** | 2.7 | 822 |
| **D4** | 6.1 | N.D |
| **D7** | 7.0 | N.D |
| **D8** | 25.4 | N.D |

**Supplementary figure 5. Anti-PvPC nanobodies broadly cross-react with PkPC, but not PfPC. a.** Representative biolayer interferometry sensorgrams of nanobody binding to PkPC and PfPC. Data are in color and 1:1 model best fit in black. **b.** Table of $K_D$ values, in nM, for the curves in a). N.D. = not determined.

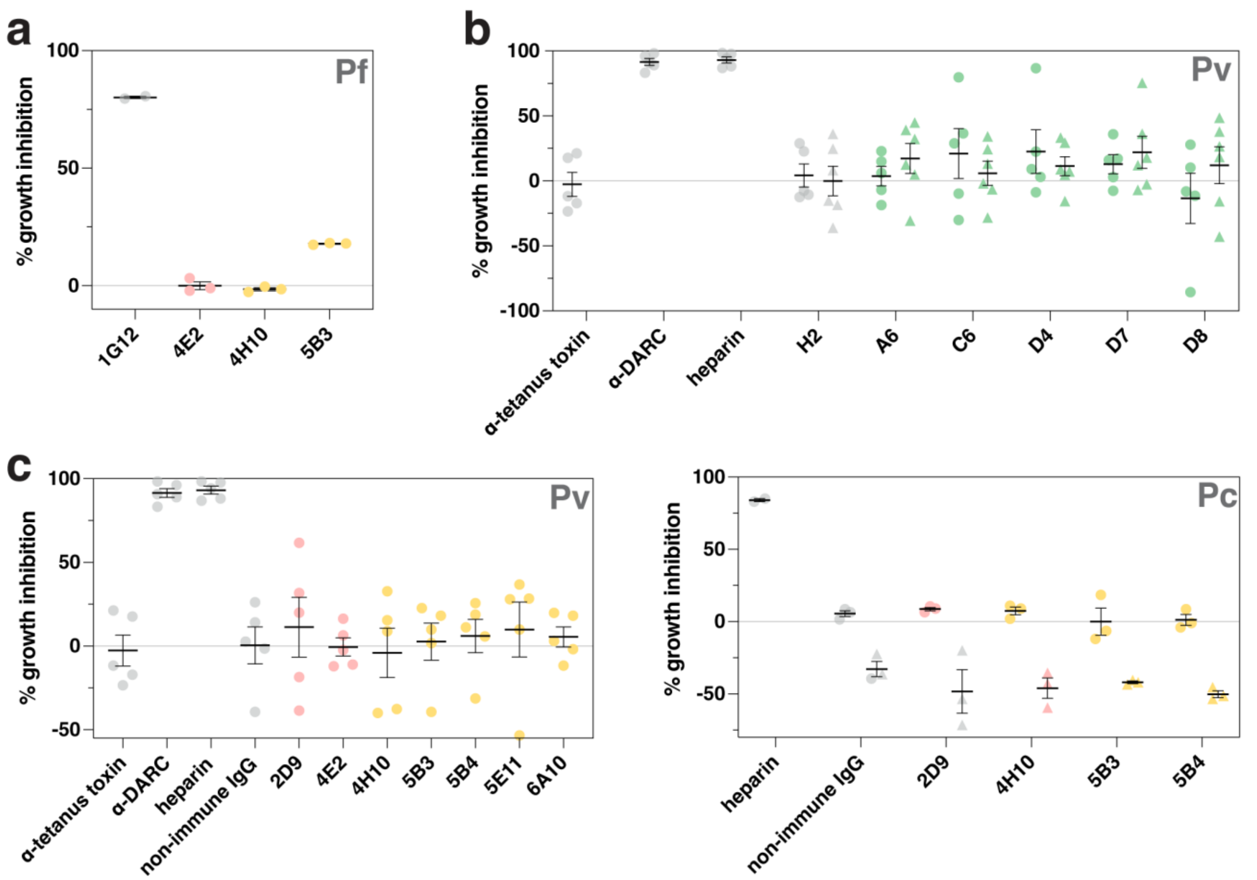

**Supplementary figure 6. Growth-inhibition assays of antibodies and nanobodies against *P. falciparum*, *P. vivax*, and *P. cynomolgi* a.** Initial screening of cross-reactive antibodies against *P. falciparum*. Antibodies were tested at a final concentration of 0.5 mg/mL for 4E2, 4H10, and 5B3, and at 1 mg/mL for 1G12 (anti-PfRipr antibody). Three biological replicates are plotted for anti-PvPC and anti-PvRipr antibodies, and two biological replicates are plotted for 1G12 **b.** *Ex vivo* growth inhibition assay of *P. vivax* parasites testing anti-PvCSS nanobodies. Circles and triangles represent 100 µg/mL and 0.5 mg/mL final concentration, respectively. Data are from five (100 µg/mL) and six (0.5 mg/mL) independent experiments. The anti-PfCSS nanobody, H2, was used as a negative control **c.** *Ex vivo* growth inhibition assay of *P. vivax* parasites. Antibodies were tested at a final concentration of 100 µg/mL. Data are from five independent experiments **d.** *P. cynomolgi* growth inhibition assay of anti-PvPC and anti-PvRipr antibodies. Three independent experiments were performed. Circles and triangles represent 125 µg/mL and 2 mg/mL final concentration, respectively. All panels show the mean and SEM.
